# Synovial fibroblast niche shapes the efficacy–safety dynamics of JAK inhibition in rheumatoid arthritis

**DOI:** 10.64898/2026.03.23.713616

**Authors:** Ana Županič, Sam G Edalat, Blaž Burja, Miriam Pauline Busch, Tadeja Kuret, Nadja Ižanc, Rahel S Zingg, Laura M Merlo Pich, Snežna Sodin-Semrl, Oliver Distler, Miranda Houtman, Caroline Ospelt, Reto Gerber, Mark D Robinson, Mojca Frank Bertoncelj

## Abstract

Synovial fibroblasts (SF) drive joint pathology in rheumatoid arthritis (RA). Difficult-to-treat RA frequently exhibits a fibroblast-rich synovial pathotype, enriched in DKK3□ and CD34□ SF, highlighting a critical therapeutic gap. Through multicohort transcriptomic analysis of synovial tissues and mechanistic *in vitro* studies, we identified SF as principal targets of Janus kinase (JAK) inhibition in RA. We demonstrated that JAK inhibitors (JAKi) can target multiple core aspects of fibroblast pathobiology – therapeutic refractoriness, cartilage destruction, and inflammation – offering a mechanistic rationale for JAKi superiority in difficult-to-treat RA. JAK1 was the dominantly expressed JAK across synovial pathotypes and SF subsets, including DKK3□ and CD34□ populations. A STAT1–interferon type I gene program was enriched in matrix-destructive PRG4□ SF, consistent with JAKi efficacy in erosive RA. In contrast, canonical IL-6 signaling predominated in IL6–expressing inflammatory CXCL12^high^ and HLA-DR+ SF, and was reproduced in cytokine-stimulated cultured SF, underscoring the autocrine nature of synovial IL-6 signaling. These data inferred a heightened JAKi sensitivity of PRG4□, CXCL12^high^, and HLA-DR+ SF subsets, informing precision therapeutic strategies. We uncovered a strong synergy between TNF and IL-6 trans-signaling, profoundly amplifying fibroblast inflammation. In high and synergistic cytokine milieu, STAT1/3 phosphorylation and IL-6 secretion persisted in SF despite tofacitinib treatment, revealing tofacitinib’s functional ceiling. This could explain reduced tofacitinib efficacy and adherence in patients with high baseline arthritis activity. Finally, inflamed SF partially uncoupled STAT3 activation from sustained JAK1 phosphorylation, limiting inflammatory output. Similar uncoupling in tofacitinib-treated SF, likely drove rapid STAT1/3 reactivation following tofacitinib washout. These data aligned with JAKi withdrawal complications and clinical recommendations for gradual JAKi tapering. Collectively, our study identifies SF as key cellular targets of JAK inhibition and delineates cytokine– and drug-driven mechanisms that may constrain the efficacy and safety profiles of JAKi in RA.

## Introduction

Rheumatoid arthritis (RA) is a destructive inflammatory arthritis affecting up to 1% of the global population (1). Despite effective disease-modifying antirheumatic drugs (DMARDs), treatment failure remains common; 30–40% of patients respond inadequately, and 5.9% to 27.5% develop difficult-to-treat RA (D2T-RA) (2, 3).

RA develops through reciprocal interactions between immune cells and joint-resident structural cells. Among these, fibroblast-like synoviocytes – synovial fibroblasts (SF) – constitute a heterogeneous, functionally specialised synovial cell population (4–7). SF activity states are further shaped by diverse mediators from structural, myeloid, and T cells, co-populating inflamed synovial tissue in RA (8). The cellular and molecular heterogeneity of synovial tissue (9), captured with synovial pathotypes (10) and cell type abundance phenotypes (11), likely contributes to interpatient variability in DMARD responses. SF may contribute to therapeutic resistance in RA (12). A stromal/fibroblast gene expression signature is enriched in synovial tissue of patients with D2T-RA (13), indicating that SF remain inadequately targeted in RA.

JAK inhibitors (JAKi) are increasingly recognised as potentially superior to other DMARDs in patients with D2T-RA (3, 14), which is likely attributable to their broad spectrum of molecular and cellular targets. By blocking signaling from numerous RA-relevant cytokines, JAKi repress multiple immune cell activities (15). At high, beyond-therapeutic concentrations, JAKi can also inhibit pathogenic activities of SF *in vitro* (reviewed in 16), indicating potential efficacy against SF. Similar to biological DMARDs, JAKi are considered for patients with poor prognostic factors who fail initial conventional DMARD therapy (17). Risk factors for adverse events, including age, smoking history, cardiovascular disease, malignancy, and thromboembolic events, must be considered when prescribing JAKi (18, 19). Abrupt JAKi discontinuation may trigger acute arthritis flares, highlighting the need for gradual drug tapering (20).

JAK inhibitors show emerging efficacy in D2T RA, where the fibroid pathotype and SF are overrepresented. We therefore aimed to delineate the cellular targets and signaling dynamics of JAK inhibition within synovial tissue and across SF subsets. Analysing multiple transcriptomics datasets, we mapped subset-specific JAK–STAT transcriptional programs in SF. Furthermore, we investigated how JAKi tofacitinib modulates SF activity across diverse cytokine environments. Our experiments mirrored highly inflamed joints as well as real-life therapy scenarios. With this design, we clarified a direct impact and caveats of tofacitinib-driven JAK inhibition on pathogenic stromal cells in RA.

## Materials and Methods

### Collection of human synovial tissue

We obtained synovial tissues from joints of RA patients undergoing joint replacement surgery at the Schulthess Clinic Zurich, Switzerland. Patients fulfilled the 2010 ACR/EULAR criteria for RA classification (21). The study was approved by the Cantonal Ethics Committee Zurich, Switzerland. All study participants signed the informed consent before enrolment. Experimental methods complied with the Helsinki Declaration. Patient characteristics and utilization of biosamples in experiments are described in **Suppl. Table 1**.

### RNA isolation from synovial tissue

For TaqMan ArrayCard experiments, we used synovial tissue, obtained from 11 RA patients during joint replacement surgery. Fragments of snap-frozen synovial tissue were powdered in a liquid nitrogen chamber. RNA was isolated using TRIzol (Life Technologies) and purified on-column, including on-column genomic DNA digestion (RNeasy Mini kit, Qiagen).

### TaqMan ArrayCard Experiments on synovial tissue

We utilised the 384-well microfluidic JAK-STAT Pathway TaqMan ArrayCards, Format 96b (Applied Biosystems) containing 6 housekeeper gene probes and 90 JAK-STAT pathway-associated gene probes. 1082 ng RNA per synovial tissue sample (n=12) was reverse transcribed (High-Capacity cDNA Reverse Transcription kit, Applied Biosystems), cDNA premixed with TaqMan Universal Master Mix II (Applied Biosystems) and loaded onto ArrayCards. Two RNA samples were loaded per ArrayCard, and gene expression was measured in duplicates per each donor. After loading, ArrayCards were centrifuged, sealed, and ran on 7900HT Real-Time PCR Instrument (Life Technologies, 40 cycles).

### Analysis of synovial tissue TaqMan ArrayCards data

TaqMan ArrayCards data on RA synovial tissue (n=12) were analysed with the comparative (ΔCt) method (22); target gene Ct values were normalised to the mean Ct of all expressed genes. Gene expression, defined as ΔCt, was plotted as box and whisker plots with median and min to max. Non-expressed genes were excluded from analysis. The created data represent Dataset 1, utilised for analysis of tissue-level expression of the JAK-STAT pathway-associated genes.

### Synovial tissue RNA-Seq Data from Pathobiology of Early Arthritis Cohort

To examine synovial tissue-level expression of JAK-STAT pathway-associated genes in an independent dataset as well as compare gene expression across synovial tissue pathotypes, we utilised the RNA-seq data from Pathobiology of Early Arthritis – PEAC Cohort (10). The data are deposited in the ArrayExpress database (Accession code E-MTAB-6141) by the owners Christopher John and Myles Lewis, Queen Mary University of London, UK, and are available on https://peac.hpc.qmul.ac.uk/ website. Normalised expression values for the JAK-STAT pathway-associated genes were extracted for 81 tissue samples classified into fibroid (n=16), diffuse myeloid (n=20) and lymphoid (n=45) pathotypes.

### ScRNA-sequencing data from synovial tissues

For analysis of gene expression across synovial cell populations and SF subsets, we utilised our scRNA-seq synovial tissue atlas of inflammatory arthritis (E-MTAB-11791, ArrayExpress, n=25 synovial biopsies, > 100000 high quality single cell profiles) (4). For Dataset 3 we focused on RA synovial biopsy data, generated from 15 synovial tissue biopsies in the atlas. The processing of scRNA-seq data was performed as described in (4). To show global (tissue-level) JAK-STAT pathway gene signatures, gene counts were summed across all cells within each sample to create pseudobulk profiles. Pseudobulk counts were subsequently normalised using sample-specific library size factors (23). The results are visualised in boxplots with each dot representing a sample. Heatmaps were used to summarize gene expression patterns across main synovial cell types / SF subsets. The mean log-normalised expression per gene and group was calculated and values were z-scored per gene across groups (24). Violin plots show the log-normalised expression stratified by main cell populations / SF subsets. Summary points (mean expression values) were overlaid as indicated in the respective figure panels of the violin plots. To compare gene expression across cell types / SF subsets while accounting for differences in abundance, we computed the log10-transformed ratio of the gene expression proportion to the cell type / SF subset proportion for each sample. For a gene *g* and group *k* (cell type / SF subset) the calculation was as follows: *log ratio(g,k) = log10 ((Cg,k/Cg,all) / (Nk/Nall))*, where *Cg,k* is a summed count for gene *g* in a cell type / SF subset *k*, *Cg,all* is a summed count for a gene *g* across all cells / SF, *Nk* is the number of cells in a cell type / SF subset and *Nall* is the number of all cells / SF. The results were visualised in boxplots with overlaid points representing individual samples. A log ratio of 0 indicates comparable expression across cell types, whereas positive or negative values indicate over– or under-expression, respectively. Dimensionality reduction for SF was performed by first computing principal components (PCA) using 3291 highly variable genes on modeled gene-wise variance (FDR < 0.05). Cells were integrated across samples using a mutual nearest-neighbour approach to reduce sample-specific batch effects. The two-dimensional UMAP embedding was generated from the batch-corrected low-dimensional space and visualised as a scatter plot where each dot represents a cell from the SF population. To improve readability of the embedding, a small number of outlier cells (2 cells) were removed based on their UMAP coordinates (cells with UMAP1 > 12 or UMAP2 < −5). Gene co-expression was assessed by comparing the distribution of logcounts for selected genes across SF subsets and visualised in ridge plots. All plots except the heatmaps were generated with ggplot2 (25). The data and code are available in Zenodo, https://doi.org/10.5281/zenodo.18395432.

### Synovial fibroblast culture

Human primary SF cultures were established from RA synovial tissue. SF were cultured in Dulbecco’s modified Eagle’s medium (DMEM F12; Life Technologies) supplemented with 10% fetal calf serum (FCS), 50□U/ml penicillin/streptomycin, 2□mM L-glutamine, 10□mM HEPES and 0.2% amphotericin B (Life Technologies). Cell cultures were negative for mycoplasma (MycoAlert mycoplasma detection kit, Lonza).

### Experiments on cultured synovial fibroblasts

SF, passages 5 to 8, were seeded in T25 flasks (400000 cells/flask, TaqMan ArrayCard experiments) or 6-well plates (100000 cells/well). For ArrayCard experiments, SF were serum-prestarved (DMEM F12, 0% FCS – starvation medium) for 6h and stimulated in starvation medium for 24h with either 0.1ng/ml TNF (Recombinant human TNF alpha, R&D Systems, 210-TA-020) + 50ng/ml sIL-6R (Recombinant human IL-6Ra, R&D Systems, 227-SR-025) or with 50ng/ml IL-6 (Recombinant human IL-6, R&D Systems, 206-IL-010/CF) + 50ng/ml sIL-6R. For other experiments, SF were serum-prestarved for 24h, and pretreated for 2h with 80nM, 180nM or 1000nM DMSO-diluted Tofacitinib Citrate [CP-690550, Lubio/Selleck Chemicals, S5001-10MM/1ML] or an equal volume of DMSO without the drug (negative vehicle control). Cell were then stimulated either with 50ng/ml sIL-6R + 50ng/ml IL-6 or with 0.1ng/ml /1ng/ml TNF ± 50ng/ml sIL-6R in a starvation medium in the presence or absence of tofacitinib. For TaqMan ArrayCards, qPCR and ELISA readouts, experiments were terminated at 24h of stimulation and for Western blot at 10min or 5h after stimulation by direct SF lysis. For washout experiments, cells were washed 3x with PBS and a fresh medium was added, with or without IL-6 + sIL-6R or tofacitinib for an additional 10min.

### RNA isolation from synovial fibroblasts

We isolated RNA by direct cell lysis in T25 flasks or 6-well plates using RNA lysis buffer, followed by the on-column RNA isolation and DNA digestion (RNeasy Mini kit, Qiagen).

### TaqMan JAK-STAT ArrayCard Experiments in synovial fibroblasts

We utilised 384-well microfluidic format 96B JAK-STAT Pathway TaqMan Array Cards (Applied Biosystems) for analysing the expression of JAK-STAT pathway-associated genes in (un)stimulated SF (n=11). cDNA synthesis and qPCR (40 cycles) were performed as described for synovial tissue. We used 285ng cDNA per port and ran measurements in duplicates; sample distribution across ArrayCards was randomised.

### Analysis of synovial fibroblast TaqMan ArrayCards data

TaqMan ArrayCard fibroblast data were analysed with the comparative (ΔCt) method (22); target gene Ct was normalised to Ct of the housekeeper gene ubiquitin C (*UBC)*. Not detected genes were excluded from analysis. Principal component analysis (PCA) was performed on ΔCt values for 72 genes across 11 donors (33 samples total, including unstimulated and two stimulated conditions) using ClustVis (https://biit.cs.ut.ee/clustvis/) (26). Rows (genes) were scaled to unit variance. Principal components (PCs) were calculated using singular value decomposition (SVD), and PC loadings were extracted for each gene to quantify their contribution to each PC. Larger magnitude loadings indicated a stronger influence of the gene on the corresponding PC. Sample-wise similarity analysis was performed in Morpheus (https://software.broadinstitute.org/morpheus/) using gene expression values (ΔCt).

Similarity between samples was calculated based on gene expression profiles using Pearson correlation and visualised as a similarity matrix. Heatmaps were generated in Morpheus, and rows and columns were hierarchically clustered using one minus the Pearson correlation coefficient with average linkage. Plots of differentially expressed genes (absolute fold change ≥ 0.5, *p* < 0.025) between stimulated and unstimulated SF were generated using GraphPad Prism Software. Gene expression was quantified as 2^ΔΔCt^, where ΔΔCt = ΔCt (stimulated SF) – ΔCt (unstimulated SF).

### cDNA synthesis and qPCR

RNA was reverse transcribed (random hexamers, MultiScribe Reverse Transcriptase, Applied Biosystems), followed by SYBR green (Roche) or TaqMan (Applied Biosystems) qPCR on 7500 or 7900HT instruments (Life Technologies). Primer and probe sequences are listed in **Suppl. Table 2.** Non template controls, dissociation curves and samples containing non-transcribed RNA were measured in parallel. Data were analysed using the comparative ΔCt method (22), normalised to *UBC*, and presented as ΔCt = Ct (Gene of interest) – Ct (*UBC*).

### Enzyme-linked immunosorbent assay

The IL-6 / IL-8 secretion was measured in cell supernatants using human IL-6/IL-8 ELISA kits (BD Biosciences) and the GloMax-Multi+Detection System (Promega). IL-6/IL-8 levels in the presence of tofacitinib were normalised to the corresponding levels in the absence of tofacitinib (0 uM) within each experimental condition. In tofacitinib experiments, data were calculated as *x-fold (s,c) = Interleukin (s,d) / Interleukin (s,0)*, where *Interleukin* denotes the cytokine (IL-6, IL-8), *s* denotes the experimental condition (unstimulated or stimulated), *d* denotes the tofacitinib concentration (0, 80, 180, 1000nM) and x-fold = 1 represents the baseline x-fold, calculated for the 0nM tofacitinib concentration within each experimental condition. This normalization allowed quantification of relative effects of tofacitinib within experimental conditions.

### Immunoblotting

For Western blotting, adherent SF were washed with PBS and lysed in 2x Laemmli buffer (BIO RAD), supplemented with cOmplete Mini Protease Inhibitor (Roche) and PhosSTOP (Roche). Lysates were boiled for 5 min before running the SDS-PAGE. After immunoblotting, nitrocellulose membranes were blocked in TBS-T buffer, containing 5% milk or 5% BSA (for phosphorylated protein detection) and incubated overnight at 4°C with primary antibodies (**Suppl. Table 3**). Membranes were then washed and incubated with HRP-conjugated anti-rabbit /-mouse IgG secondary antibodies (Jackson ImmunoResearch, 1:10000) in TBS-T, containing 5% milk or BSA. Protein bands were detected and quantified using chemiluminescence method (WesternBright ECL HRP substrate, Advantsta) on Fusion FX instrument (Vilber). Original Western blot images are provided in **Suppl. Figs. 3-6**.

### Statistical analysis

Data were analysed using GraphPad Prism v.8.0. Data distribution was assessed with the Anderson-Darling, D’Agostino and Pearson, Shapiro-Wilk and Kolmogorov-Smirnov tests. For paired two-group comparisons, a two-tailed paired t-test (normally-distributed data) or the Wilcoxon matched-pairs signed-rank test (not-normally-distributed data) was used. For multigroup comparisons of paired data, repeated-measures ANOVA (normally-distributed data) or the Friedman test (not-normally-distributed data) were performed. p-values were adjusted for multiple comparisons using Tukey’s / Šídák’s or Dunn’s tests, respectively. A one-sample t-test or Wilcoxon test (for not-normally-distributed data) were used to assess whether the mean or median, respectively, differed from a theoretical value (1 or 0), with Bonferroni correction for multiple comparisons. Unless otherwise indicated, p-values < 0.05 were considered statistically significant.

## Results

### JAK1 is the highest expressed Janus kinase gene in RA synovial tissue

We characterised the global JAK–STAT gene signature in RA synovial tissue using three independent gene expression datasets. The key features of the JAK-STAT pathway are presented in **Fig. 1a**. Characteristics of the used gene expression datasets are summarised in **Fig. 1b**. Dataset 1 comprises in-house gene expression data from RA synovial tissue (**Fig. 1c**); RA patients had diverse clinical and treatment histories (**Suppl. Table 1**). Dataset 2 represents published synovial RNA-seq data from PEAC cohort (10); in this dataset we explored global (**Fig. 1d**) and pathotype-enriched JAK–STAT pathway gene signatures. Dataset 3 includes in-house scRNA-seq data (E-MTAB-11791) from RA patients with heterogeneous clinical and therapeutic backgrounds (4). These data are part of our published single cell synovial tissue atlas in inflammatory arthritis (4). In Dataset 3, we studied the tissue-level (**Fig. 1e**) and cell type-specific JAK–STAT pathway gene signatures.

**Figure 1.**
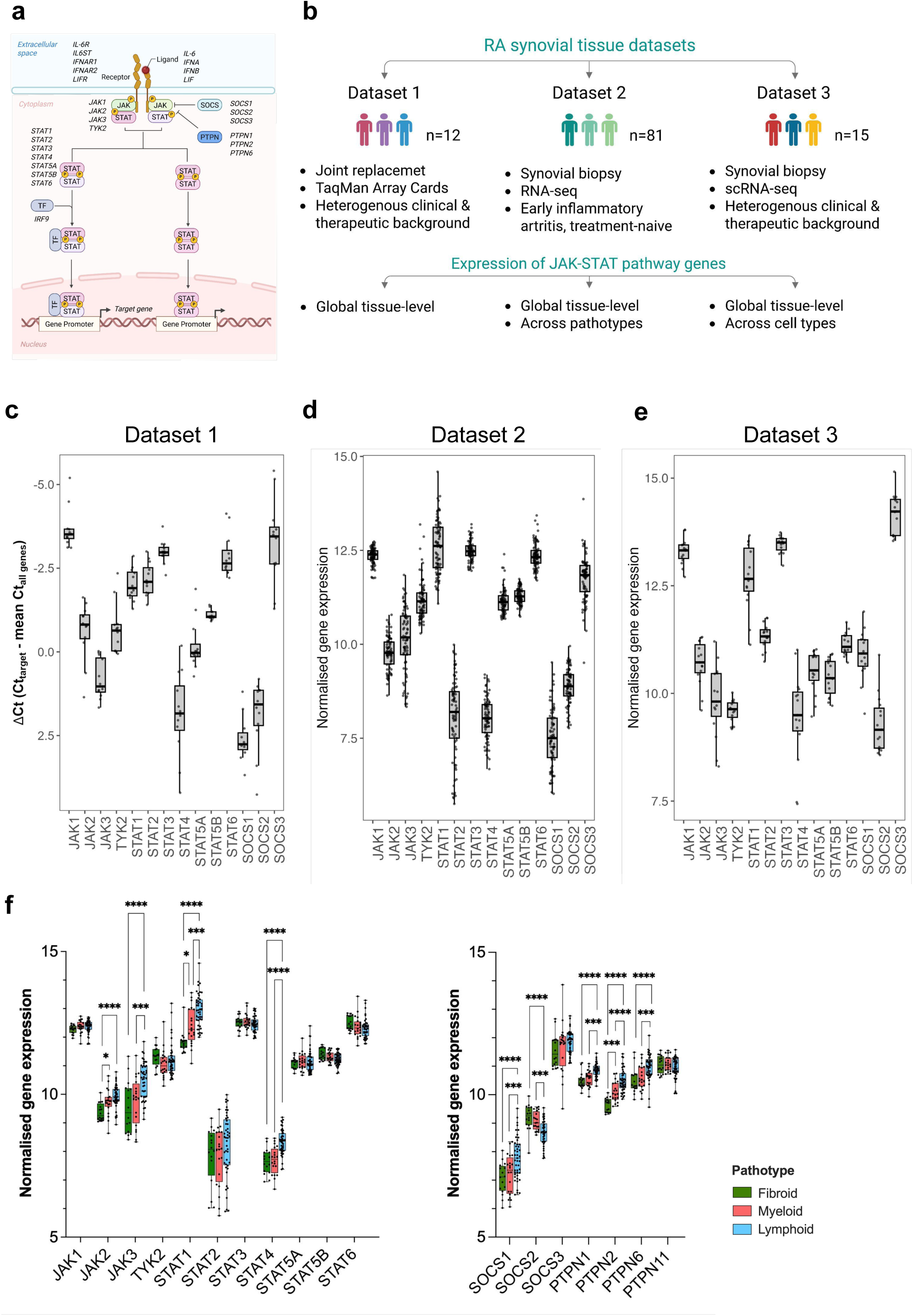
RA synovial tissue and synovial pathotypes display shared and unique patterns in JAK-STAT pathway-associated genes. **a)** A scheme denoting transcriptional signature of JAK-STAT pathway-associated genes. These genes include canonical JAK and STAT genes, pathway regulators, ligands, receptors and transcriptional factors. Created in BioRender, https://BioRender.com/90tyzjw, 2026. **b)** Characteristics of RA datasets utilised in analysis of the JAK-STAT pathway transcriptional signature in synovial tissue. Created with BioRender. N = 108 synovial tissue samples from three independent datasets. The datasets include an in-house Dataset 1, Dataset 2 – PEAC RNA-seq Cohort (10) and RA scRNA-seq synovial tissue Dataset 3. Dataset 3 comprises scRNA-seq data from 15 RA synovial tissues, extracted from our scRNA-seq synovial tissue atlas in inflammatory arthritis (BioStudies Database, ArrayExpress, accession number E-MTAB-11791, (4)). **c)** The expression of JAK, STAT and SOCS genes in Dataset 1. Gene expression, measured using the JAK-STAT Pathway TaqMan ArrayCards, was quantified as ΔCt. Data are presented as box and whiskers plot, with median ΔCt and min to max. **d)** Normalised expression of JAK-STAT pathway-associated genes in 81 synovial tissues from Dataset 2. RNA-seq data was extracted from https://peac.hpc.qmul.ac.uk/ (10). **e)** Pseudobulk normalised expression of JAK-STAT pathway-associated genes across synovial tissue samples from scRNA-seq Dataset 3 (4). **f)** Normalised expression of JAK-STAT pathway-associated genes across fibroid (n=16), diffuse myeloid (n=20) and lymphoid (n=45) synovial tissue pathotypes from Dataset 2; RNA-seq data was extracted from https://peac.hpc.qmul.ac.uk/ (10).

Tissue-level analyses demonstrated that *JAK1*, *STAT3*, and *SOCS3* were consistently highly expressed, and *JAK3* and *STAT4* lowly expressed (**Fig. 1c-e**) in RA synovial tissue. The dominant synovial tissue expression of JAK1 is consistent with the observed clinical benefit of JAK1 inhibition in RA (27).

### Shared and unique JAK–STAT pathway gene signatures underpin synovial pathotypes

Transcriptional heterogeneity at the tissue level reflects, among other factors, the underlying variation in cellular composition. Synovial pathotype (10) and CTAP (11) classifications capture cellular variability in RA synovial tissue and correlate with DMARD responses (13, 28, 29). Accordingly, we investigated how synovial cell types contributed to the tissue-level JAK–STAT transcriptional signature, utilising a two-step approach. Specifically, we compared the expression of target genes across synovial tissue pathotypes in Dataset 2 (10) as well as investigated the expression of the JAK-STAT pathway genes across synovial cell types in Dataset 3.

Pathotype classification categorizes synovial tissue as diffuse-myeloid, lymphoid, or immune cell–sparse, fibroblast-rich/fibroid (10). We revealed comparable expression of *JAK1*, *TYK2*, and most *STAT* genes across synovial tissue pathotypes (**Fig. 1f**). By contrast, JAK2, JAK3, STAT1 and STAT4 were enriched in the lymphoid pathotype and underrepresented in the fibroid pathotype (**Fig. 1f**). Genes encoding the JAK-STAT pathway regulators were either comparably expressed across pathotypes (e.g. *SOCS3, PTPN11*) or pathotype-enriched (*SOCS1/2, PTPN1/2/6*; **Fig 1f**). The observed differences, while statistically significant, were small in magnitude.

### Synovial cells exhibit overlapping and distinct JAK–STAT pathway gene expression programs

To explore the JAK–STAT pathway gene expression across synovial cell types, we analysed 62,470 single cell transcriptomes from scRNA-seq Dataset 3. Synovial cells spanned 3 structural and 8 leukocyte cell populations (4). We identified three principal gene expression modules, represented by distinct JAK–STAT pathway genes (**Fig. 2a**); a structural cell-enriched module, mixed myeloid–B cell module, and a T cell module (**Fig. 2a**). These data inferred that synovial myeloid, lymphoid and structural cells may preferentially utilise different JAK-STAT signalling trajectories. *JAK1*, *STAT3*, and *SOCS3* were expressed in all cell populations, with levels varying by cell type (**Fig. 2b**). For example, *JAK1* showed higher expression in lymphocytes, dendritic cells, and endothelial cells (**Fig. 2a, b**). *SOCS3* was most enriched in synovial structural cells (**Fig. 2a, b**). By contrast, *PTPN1* and *PTPN2* transcripts were abundant in dendritic cells and B cells/plasmablasts (**Fig. 2a, Suppl. Fig. 1a**). Other JAK, STAT, and pathway regulator genes were detected at low levels in the scRNA-seq data and commonly exhibited cell type–dominated expression (**Fig. 2b; Suppl. Fig. 1a**).

**Figure 2.**
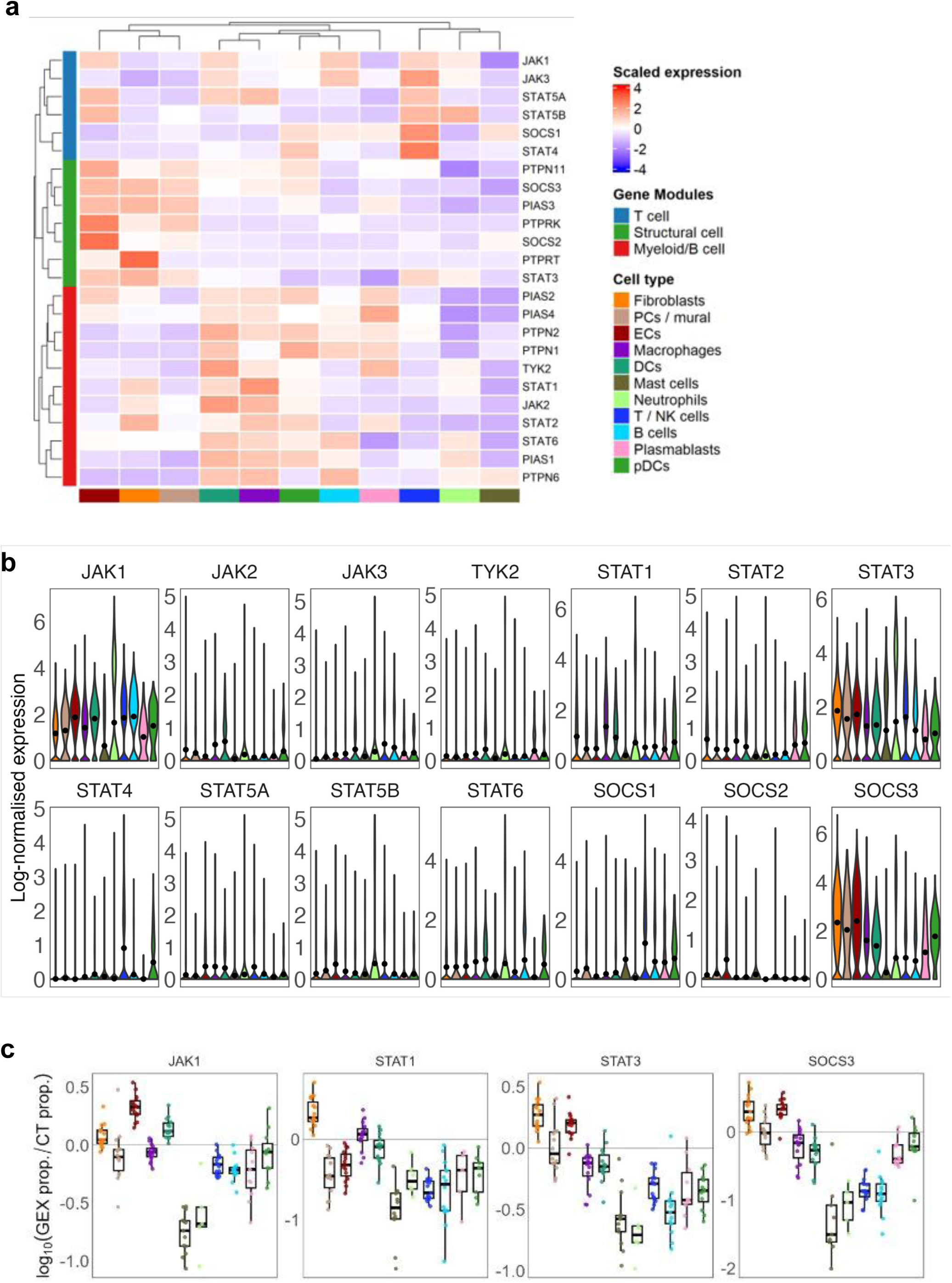
Synovial cell types display shared and unique JAK-STAT pathway gene signatures a-c) Expression of the. JAK-STAT pathway-associated genes across synovial structural, myeloid and lymphoid cell populations in scRNA-seq. For Dataset 3, we extracted RA scRNA-seq from our synovial scRNA-seq tissue atlas in inflammatory arthritis (BioStudies Database, ArrayExpress, accession number E-MTAB-11791, (4)). **a)** The heatmap shows the mean log normalised expression per gene and group, expression values are z scored per gene across groups. **b)** The violin plots show the log normalised expression of JAK-STAT pathway genes across principal synovial cell types. **c)** Contribution of synovial cell types to total synovial JAK-STAT pathway transcriptional signature, calculated as log10-transformed ratio between gene expression proportion (GEX prop.) and cell type proportion (CT prop.) per sample. A log10 ratio of 0 indicates comparable expression across cell types, whereas positive and negative values indicate over– and under-expressed genes, respectively, in a given cell type versus all cell types. PCs: pericytes, ECs: endothelial cells, DCs: dendritic cells, NK: natural killer, pDCs: plasmacytoid dendritic cells.

Cell type-dominated gene expression frequently coincided with pathotype-enriched transcriptional signatures. Specifically, JAK3, STAT4 and SOCS1 transcripts were enriched in lymphocytes (**Fig. 2a, b**), and SOCS2 in endothelial cells (**Fig. 2a, b**). This expression pattern aligned with their overrepresentation in lymphoid and fibroid pathotypes (Fig. 1f), respectively. *PTPN1/2/6* showed highest expression in lymphocytes and dendritic cells (**Fig. 2a, b, Suppl. Fig. 1a**), aligning with their overrepresentation in lymphoid pathotype (**Fig. 1f**).

Beyond transcript quantity, cell type abundance shapes the tissue-level gene expression. Accordingly, we integrated transcript quantities and cell type abundances and quantified cell type contributions to the global JAK–STAT pathway transcriptional signature in synovial tissue (**Fig. 2c, Suppl. Fig. 1b**). We focused on JAK-STAT pathway-associated transcripts, readily detected in scRNA-seq Dataset 3 (**Fig. 2b, Suppl. Fig. 1a**). SF and endothelial cells contributed stronger to total synovial *JAK1, STAT3 and SOCS3* expression, whereas SF and macrophages added to *STAT1* and *PTPN2* expression (**Fig. 2c, Suppl. Fig. 1b**). Plasmablasts/B cells and macrophages contributed to *PTPN1/2* and dendritic cells to *JAK1* and *PTPN1/2* expression (**Fig. 2c, Suppl. Fig. 1b**).

In summary, both immune and structural cells contributed to synovial expression of the JAK-STAT pathway-associated genes, exhibiting shared and cell type-unique expression patterns.

### Synovial fibroblasts are prominent contributors to the synovial JAK–STAT pathway gene expression in RA

We identified SF as prominent contributors to the JAK–STAT pathway gene signature in RA synovial tissue (**Fig. 2**). SF comprise functionally distinct cell subsets (4–7). DKK3+ and CD34+ SF subsets appear expanded in RA patients who fail to respond to biological DMARDs (13, 30). Accordingly, we examined the expression of the JAK-STAT pathway genes in 19,907 single COL1A1+ SF from the Dataset 3 (4).

We first assessed the global JAK–STAT pathway gene expression signature in total COL1A1□ RA SF population. Therefore, we aggregated single fibroblast transcriptomes into a pseudobulk expression profile. RA SF displayed a similar JAK-STAT pathway gene expression pattern (**Fig. 3a**) as observed in RA synovial tissue as a whole (**Fig. 1e**).

**Figure 3.**
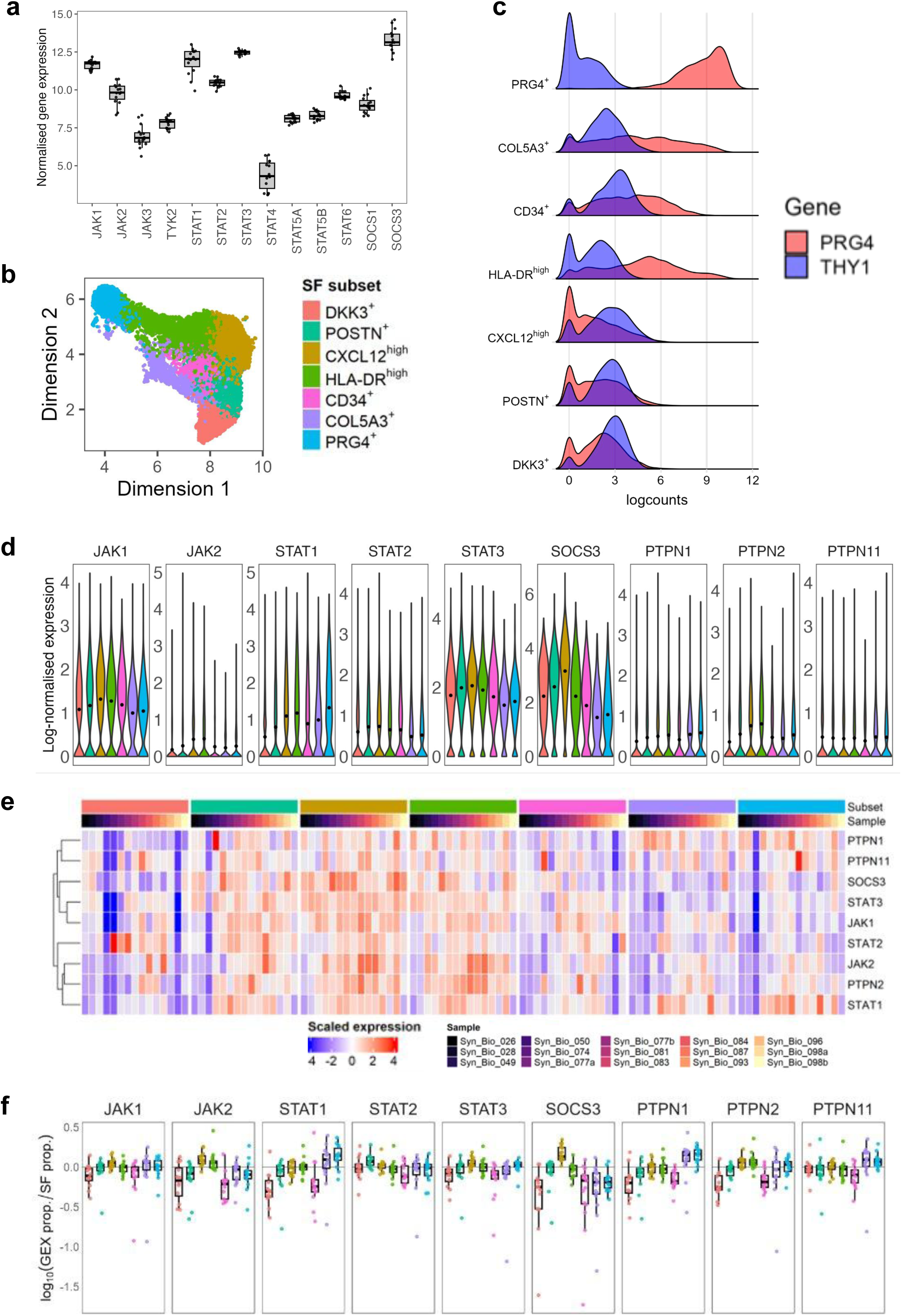
JAK-STAT pathway gene features infer a broad targeting potential of JAK inhibitors across synovial fibroblast subsets. **a-f**) Analysis of JAK STAT pathway transcriptional signature in *in silico* sorted COL1A1+ rheumatoid arthritis synovial fibroblasts (SF, n=21433) from scRNA-seq Dataset 3. For Dataset 3, we extracted RA scRNA-seq from our synovial scRNA-seq tissue atlas in inflammatory arthritis (BioStudies Database, ArrayExpress, accession number E-MTAB-11791, (4)). **a)** Normalised pseudobulk gene expression of JAK, STAT and SOCS genes in SF aggregated per sample. **b)** A UMAP plot showing clustering of COL1A1+ SF into seven distinct subsets. **c)** Ridge plots displaying the co-expression of PRG4 and THY1 genes across SF subsets, defining PRG4^high^ THY1^low^ lining SF and PRG4^low/med^ THY1+ sublining SF subsets. **d)** Violin plots showing the expression of canonical JAK and STAT genes as well as genes encoding the pathway regulators across SF subsets. **e)** Heatmap showing the expression of canonical JAK and STAT genes, and genes encoding the pathway regulators. Mean log normalised expression per gene is shown with values z-scored across groups. **f)** Contribution of SF subsets to total fibroblast JAK-STAT pathway gene signature. The contribution is calculated as log10-transformed ratio between gene expression proportion (GEX prop.) and SF subset proportion (SF prop.) per sample. A log10 ratio of 0 indicates comparable expression across cell types, whereas positive and negative values indicate over– and under-expressed genes, respectively, in a given SF subsets versus all SF.

We next delineated the JAK-STAT pathway transcriptional profiles in individual SF subsets. We focused on genes detectable in fibroblast scRNA-seq data (**Fig. 2b**). SF segregated into seven transcriptionally distinct subsets (**Fig. 3b**), spanning PRG4^high^ THY1^low^ lining and PRG4^low/med^ THY1+ sublining SF (**Fig. 3c**). *JAK1*, *STAT3*, and *SOCS3* genes were expressed across SF subsets (**Fig. 3d**), including therapeutic resistance-linked CD34+ and DKK3+ SF subsets. Overall, the JAK-STAT pathway gene signature was more pronounced in CXCL12^high^ and HLA-DR+ SF, and weaker in DKK3+, COL5A3+ and PRG4+ SF (**Fig. 3e**). CXCL12^high^ SF added strongest to *JAK2* and *SOCS3* expression, and PRG4+ SF to *STAT1* and *PTPN1* expression (**Fig. 3f**). By contrast, DKK3+ SF and CD34+ SF contributed less to the expression of several JAK-STAT pathway genes (**Fig. 3f**).

Taken together, these data suggested that CXCL12^high^, HLA-DR+ SF and PRG4+ subsets are likely to be particularly sensitive to JAK inhibition, with other subsets comparatively less so. Moreover, distinct SF subsets might preferentially engage discrete JAK–STAT signalling programmes; for example, lining PRG4□ SF appeared biased toward STAT1-driven signalling.

### Autocrine IL-6 production may amplify the *JAK1*–*STAT3*–*SOCS3* gene expression network in CXCL12^high^ and HLA-DR+ synovial fibroblasts

We showed that JAK-STAT pathway gene signatures varied across SF subsets. Thus, we next assessed the JAK-STAT pathway signalling networks in SF (**Fig. 4a**). We focused on cytokine-driven JAK-STAT signalling networks, relevant for SF and arthritis pathology. Networks included genes encoding ligand-receptor pairs, JAKs, STATs, pathway regulators, transcription factors and downstream target genes (**Fig. 4a**).

**Figure 4.**
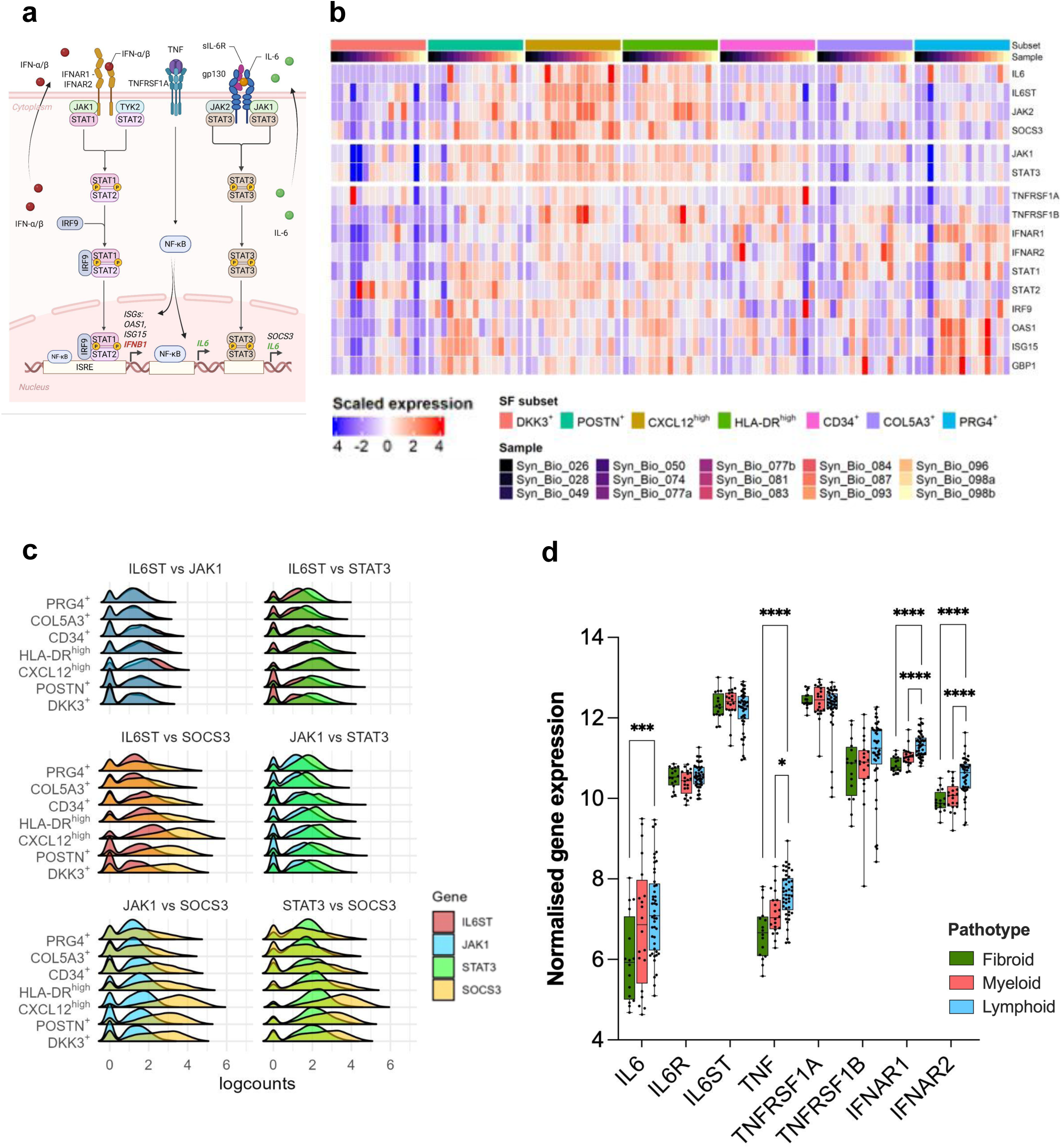
Cytokine-driven JAK-STAT signalling networks are distinctly enriched across synovial fibroblast subsets and tissue pathotypes in rheumatoid arthritis. **a**) A scheme denoting transcriptional signature of IL-6 and type I interferon-driven JAK-STAT signaling networks. Indirect signalling through TNF is depicted. The signalling networks include canonical pathway components, transcriptional factors and target downstream genes. Created in BioRender, https://BioRender.com/bc6pcc9, 2026. **b-d)** Analysis of JAK STAT pathway transcriptional signature in *in silico* sorted COL1A1+ synovial fibroblasts (SF) from scRNA-seq Dataset 3 (n=21433 single SF profiles). For Dataset 3, we extracted RA scRNA-seq from our synovial scRNA-seq tissue atlas in inflammatory arthritis (BioStudies Database, ArrayExpress, accession number E-MTAB-11791, (4)). **b)** Heatmap showing the expression of genes, encoding the core components of the IL-6– as well as type I interferon-driven JAK-STAT signalling pathways across SF subsets. TNF receptor genes are also included given critical roles of TNF in indirect JAK-STAT pathway activation. Mean log normalised expression per gene is shown, with values z-scored across groups. **c)** Ridge plots displaying co-expression of genes encoding the IL-6 associated JAK-STAT pathway components across SF subsets. **d)** Normalised expression of genes, encoding the JAK-STAT pathway-linked cytokines and their receptors across fibroid (n=16), diffuse myeloid (n=20) and lymphoid (n=45) synovial tissue pathotypes from Dataset 2. Dataset 2 RNA-seq data was extracted from https://peac.hpc.qmul.ac.uk/ (10).

We investigated the *IL6* – *IL6ST* – *JAK1/2* – *STAT3* – *SOCS3* gene network (**Fig. 4a**), encoding the core components of the IL-6–induced JAK–STAT pathway (31, 32). This network was overrepresented in the IL-6 expressing CXCL12^high^ and HLA-DR+ SF (**Fig. 4b, c**), whereas underrepresented in DKK3+ SF, COL5A3+ SF and PRG4+ SF. These findings inferred that autocrinally-produced IL-6 likely drives the JAK–STAT signalling in CXCL12^high^ and HLA-DR+ SF (**Fig. 4b, c**), which could exhibit heightened sensitivity to JAK inhibition in RA.

### Lining PRG4+ synovial fibroblasts might favor direct interferon type I signalling

In addition to IL-6, many other cytokines can activate JAK-STAT signaling (**Fig. 4a**) in RA. Type I interferons – particularly IFNβ – induced JAK-STAT signaling in SF; both directly (33) and indirectly – via TNF-induced IFNβ production (34). Expanding on these findings, we examined type I interferon-linked gene networks across SF subsets.

The TNF-type I interferon-linked gene module was lowest expressed in DKK3+ SF (**Fig. 4b**). PRG4+ SF expressed the highest level of interferon response genes (*ISG15*, *OAS1*, *GBP1*), consistent with their pronounced *IFNAR1*, *STAT1* and *IRF9* gene expression (**Fig. 4b**). These cells were low in TNF receptor gene signature, possibly indicating a preference for a direct type I interferon signalling.

Together, cytokine-receptor-signalling gene networks supported the differential engagement of JAK–STAT signalling trajectories across SF subsets.

### Expression of JAK–activating cytokines and receptors varies across synovial tissue pathotypes

The production of JAK-STAT pathway-activating cytokines – including IL□6, type I interferons, and TNF – differs among structural, myeloid, and lymphoid synovial cell populations. These cell populations themselves exhibit patient– and pathotype-specific enrichment patterns (4, 9, 11). Accordingly, we analysed the expression patterns of cytokine-receptor pairs across synovial pathotypes utilizing the Dataset 2 (10).

TNF, IL-6, and IFNAR1/2 transcripts were significantly elevated in diffuse myeloid and/or lymphoid pathotypes (**Fig.**□**4d**), although the differences were moderate in magnitude. In contrast, expression of type I interferon (**Suppl. Fig.**□**1c**) and TNF/IL□6 receptor genes (**Fig.**□**4d**) was similar across pathotypes. These data suggested that variation in cell composition may distinctly shape local synovial cytokine milieu in RA synovial tissue (4,□11). However, how moderate transcriptional differences – detected at the tissue-level – translate into local cytokine variability at the cell-cell level, remains to be determined.

### STAT phosphorylation patterns differ in IL-6– and TNF–stimulated fibroblasts

To investigate how distinct synovial cytokine milieus regulate JAK–STAT signalling in SF, we designed a panel of *in vitro* experiments in cultured RA SF. The JAK–STAT gene expression signature in cultured SF (**Suppl. Fig. 2a**) largely mirrored the fibroblast JAK-STAT pathway gene profiles in scRNA-seq Dataset 3 (**Fig. 3a**).

We stimulated SF with IL-6 and/or TNF, with or without sIL-6R. Although membrane IL-6R expression is limited in SF, gp130 (*IL6ST*) abundance enables IL-6 trans-signalling (35) within the sIL-6R-enriched RA joints (36). Early (10min) and delayed (5h) JAK-STAT pathway activation was assessed by quantifying tyrosine phosphorylation of JAK1 (Tyr1022), STAT1 (Tyr701), and STAT3 (Tyr705). These tyrosines represent characteristic phospho-sites in type I interferon and gp130-linked JAK-STAT signalling (37). IL-6 + sIL-6R robustly induced pJAK1, pSTAT1, and pSTAT3 at both timepoints (**Fig. 5a, b, Fig. 7a, b**). The IL-6 + sIL-6R-induced change in the pJAK1/JAK1 ratio was comparable at 10min and 5h (**Fig. 5c**), indicating sustained JAK1 activity. In contrast, pSTAT1/STAT1 and pSTAT3/STAT3 induction were significantly lower at 5h compared to 10min (**Fig. 5c**). These results may reflect elevated basal pSTAT1 levels at 5h (**Fig. 7b**). Moreover, inflamed SF could increase the expression of negative pathway regulators. Accumulated negative regulators could decouple STAT3 signalling from sustained JAK1 activation in IL-6–rich environments.

**Figure 5.**
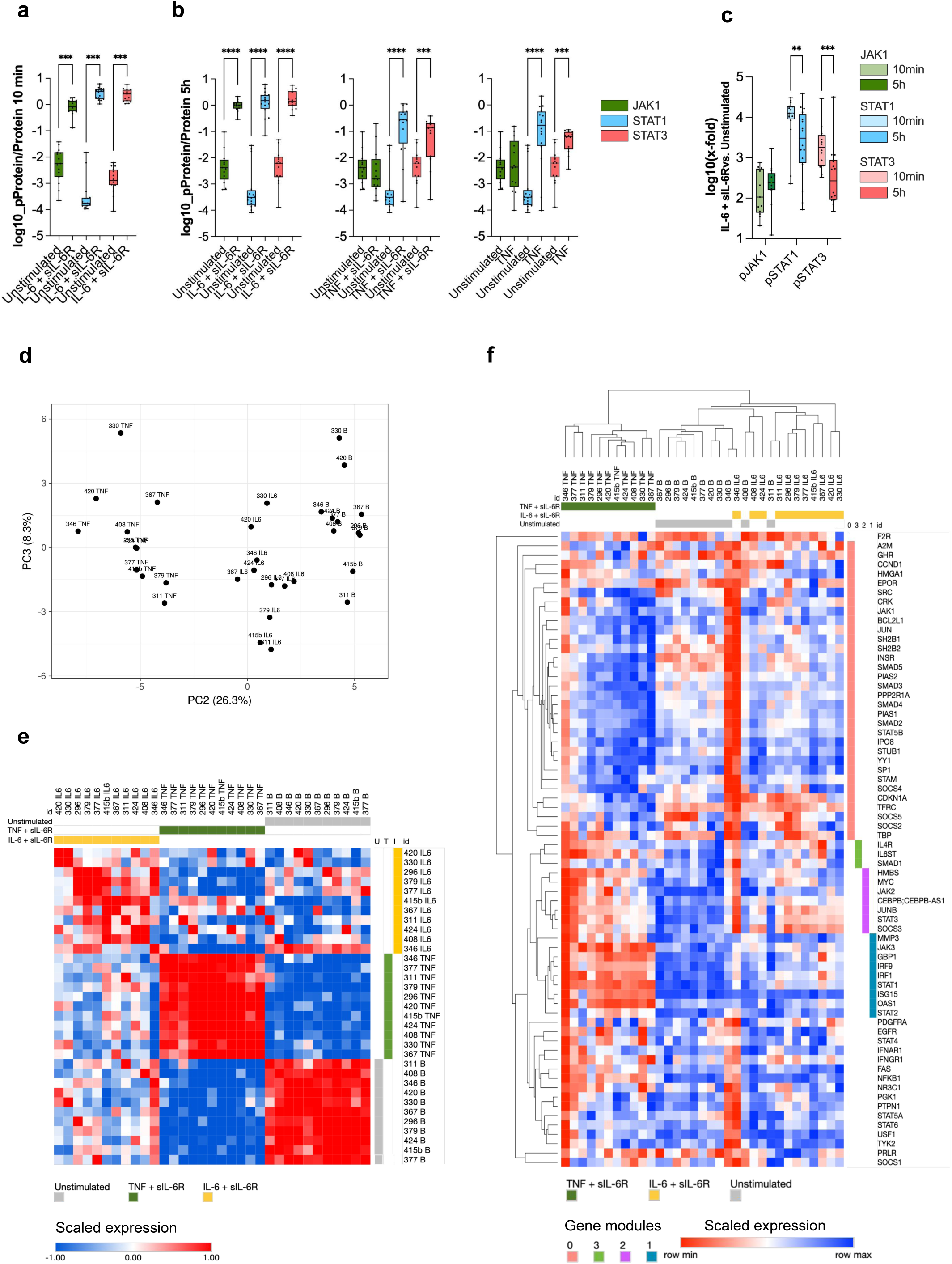
Diverse cytokine environments activate JAK-STAT pathway in rheumatoid arthritis synovial fibroblasts. **a-f**) *In vitro* experiments. We stimulated cultured rheumatoid arthritis (RA) synovial fibroblasts (SF) with 50ng/ml interleukin 6 (IL-6) or 1ng/ml tumor necrosis factor (TNF) with or without 50ng/ml soluble IL-6 receptor (sIL-6R). **a, b)** Basal and cytokine-induced phosphorylation of JAK1 and STAT1/3 proteins in SF stimulated with **a)** IL-6 + sIL-6R for 10min, n=15-16 biological replicates or **b)** IL-6 + sIL-6R, TNF + sIL-6R or TNF for 5h, n=13-16 biological replicates. We quantified the data as a ratio of phosphorylated versus total protein (pProtein/Protein), using optical density values of chemiluminescently-detected immunoblotted proteins. pProtein/Protein values were log10 transformed. Representative immunoblot images are shown in Fig. 7, and original images in **Suppl. Figs. 3, 4**. **c)** Comparison of early (10min) versus late (5h) induction of pJAK1 and pSTAT1/3 in SF, calculated as x-fold = pProtein/Protein (stimulated) / pProtein/Protein (unstimulated) per time point; x-fold values were log10 transformed. **a-c)** We present experimental data using box and whisker plots with median and min to max. We compared paired experimental groups using paired t test or Wilcoxon matched-pairs signed rank test. *p<0.05, **p<0.01, *** p<0.001, ****p<0.0001. **d-f)** Expression of JAK-STAT pathway-associated genes, probed on TaqMan ArrayCards, in unstimulated and stimulated (IL-6/TNF + sIL-6R) RA SF; n=11 biological replicates per experimental condition. We quantified gene expression as ΔCt; 72 genes were reliably detected across n = 33 experimental conditions. Not detected genes were excluded from analysis. **d)** Principal component (PC) analysis of ΔCt values using ClustVis. PC2 and PC3 separated samples according to stimulus and donor, respectively. PC1 was dominated by a single outlier sample (**Suppl.** Fig. 2b). **e)** Similarity matrix, computed for columns (samples) using Pearson correlation in Morpheus. **f)** Heatmap generated in Morpheus using ΔCt gene expression values, with hierarchical clustering of genes and experimental conditions. Rows and columns were clustered using 1 − Pearson correlation with average linkage.

**Figure 6.**
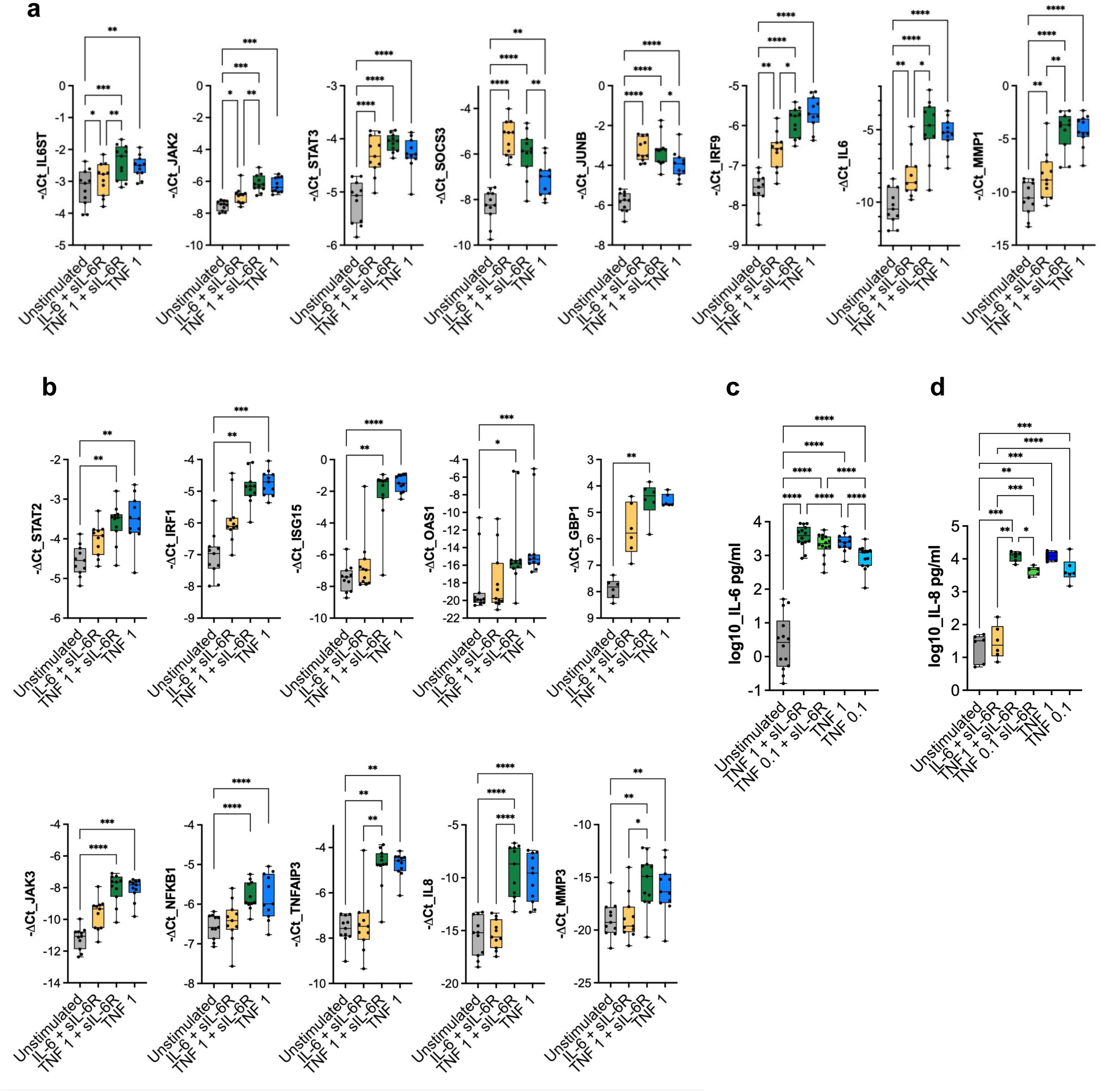
Cytokine microenvironments drive inflammatory synergies in rheumatoid arthritis synovial fibroblasts. **a, b**) *In vitro* experiments. We stimulated cultured rheumatoid arthritis (RA) synovial fibroblasts (SF) with 50ng/ml interleukin 6 (IL-6) or 1ng/ml tumor necrosis factor (TNF) ± 50ng/ml soluble IL-6 receptor (sIL-6R) for 24h. N = 6-11 biological replicates per experimental condition. Gene expression was quantified using qPCR and calculated as ΔCt. We show ΔCt using box and whisker plots with median and min to max. To evaluate differences among preselected biologically meaningful pairs of experimental conditions (unstimulated, stimulated), we used RM one-way ANOVA with Šídák’s multiple comparisons test or Friedman test with Dunn’s multiple comparisons test. **a)** Genes upregulated in the presence of both IL-6 + sIL-6R and TNF ± sIL-6R. **b)** Genes selectively upregulated with TNF ± sIL-6R. **c, d)** *In vitro* experiments. We stimulated cultured RA SF with 50ng/ml IL-6, 0.1 or 1ng/ml TNF in the presence or absence of 50ng/ml sIL-6R. Secreted **c)** IL-6 (n=12-14 biological replicates) and **d)** IL-8 (n=5-6 biological replicates) proteins were analysed 24h after stimulation using ELISA. Protein concentration values (pg/ml) were log10-normalised. Data are presented using box and whiskers plots with median and min to max. To evaluate differences among preselected biologically meaningful pairs of experimental conditions (unstimulated, stimulated), we used mixed-effects analysis with Šídák’s multiple comparisons test. *p<0.05, **p<0.01, *** p<0.001, ****p<0.0001.

**Figure 7.**
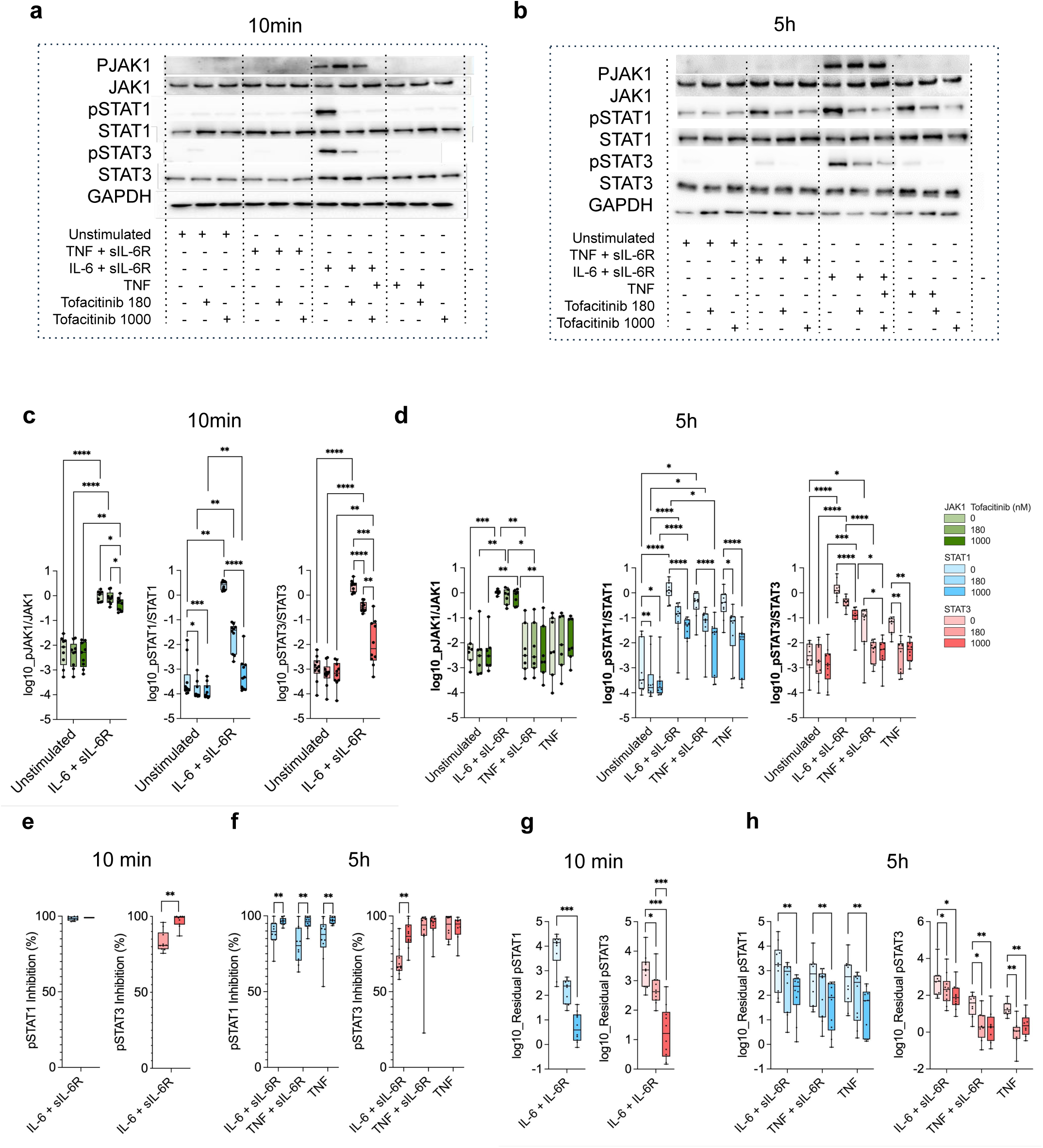
Tofacitinib therapy exhibits ceiling effects in rheumatoid arthritis synovial fibroblasts in the highly inflamed milieu. **a-h**) *In vitro* experiments. We stimulated cultured rheumatoid arthritis synovial fibroblasts (SF) with 50ng/ml interleukin 6 (IL-6) or 1ng/ml tumor necrosis factor (TNF) in the presence or absence of 50ng/ml soluble IL-6 receptor (sIL-6R) for 10min or 5h across tofacitinib concentrations (0, 180, 1000nM). Cells were pretreated with tofacitinib for 2h. **a, b)** Representative immunoblot images of (un)phosphorylated JAK1 and STAT1/3 proteins as well as GAPDH (loading control) in unstimulated SF and SF, stimulated with **a)** IL-6 and sIL-6R for 10min. N = 9-10 biological replicates or **b)** IL-6 + sIL-6R or TNF ± sIL-6R for 5h, n=7-10 biological replicates. Original immunoblot images are shown in **Suppl. Figs. 3, 4**. **c, d**) We quantified immunoblot data as a ratio of phosphorylated versus total protein, using optical density values of chemiluminesce-detected immunoblotted proteins. Data are presented using box and whisker plots with median and min to max. To compare the effects of different experimental conditions (stimulated, unstimulated) at individual tofacitinib concentrations, we used **c)** paired t-test / Wilcoxon matched-pairs signed-rank test or **d)** RM one-way ANOVA with Šídák’s multiple-comparisons test / Friedman test with Dunn’s multiple-comparisons test. Biologically meaningful pairwise comparisons were performed. To evaluate the effects of different tofacitinib concentrations (0, 180, 1000um) within each experimental condition, we performed RM one-way ANOVA with Tukey’s multiple-comparisons test or Friedman test with Dunn’s multiple-comparisons test. **e, f)** Percent inhibition of pJAK1 and pSTAT1/3, normalised to total JAK1 and STAT1/3, respectively, in IL-6 + sIL-6R-treated SF at **e)** 10min and **f)** 5h time points. We compared tofacitinib concentration effects using paired t-test or Wilcoxon matched-pairs signed-rank test. **g, h)** Residual pJAK1 and pSTAT1/3 phosphorylation, normalised to total JAK1 and STAT1/3, respectively, in tofacitinib-treated cytokine-stimulated SF. *Log10 [x-fold(s,d)] = log10 [pProtein/Protein(s,d) / pProtein/Protein*(*0,0*)], where *s* denotes stimulus (IL-6 + sIL-6R, TNF + sIL-6R, TNF), *d* represents tofacitinib concentration (0, 180, 1000nM) and *baseline log10 [x-fold(0,0] equals 0* in unstimulated SF at 0 uM tofacitinib. Data are shown using box and whisker plots with median and min to max. To compare tofacitinib concentration effects within individual stimuli, we performed RM one-way ANOVA with Tukey’s multiple-comparisons test or the Friedman test with Dunn’s multiple-comparisons test. To compare log10 [x-fold(s,d)] values to baseline log10[x-fold] we used a one-sample t-test or Wilcoxon test. p-value was adjusted for multiple comparisons using the Bonferroni test. *p<0.05, **p<0.01, *** p<0.001, ****p<0.0001.

TNF ± sIL-6R induced a distinct pattern of JAK–STAT pathway activation. pJAK1 was not significantly induced across experimental timepoints and pSTAT1/3 were increased at the 5h time point (**Fig. 5b**, **Fig. 7b**). These results indicated a delayed JAK–STAT signalling, possibly driven by the autocrinally produced IL-6 and/or type I interferons (34).

### TNF and IL-6 drive distinct transcriptional programs in synovial fibroblasts

To further dissect JAK-STAT pathway activation in SF, we measured gene expression changes in cytokine-stimulated SF, using JAK-STAT pathway TaqMan ArrayCards. Seventy-two of the 90 genes probed on the ArrayCards were reliably detected in SF. Principal component analysis of the 72 expressed genes indicated that inflammatory stimulus (PC2) and donor identity (PC3) accounted for 26.3% and 8.3% variation in gene expression (**Fig. 5d**). PC1 was dominated by samples from a single outlier donor (**Suppl. Fig. 2b**). PC loadings analysis demonstrated that interferon response genes, subset of JAK-STAT genes and inflammatory genes contributed strongest to stimulus-specific variability (**Suppl. Table 4**). Meanwhile, the expression of several IL-6 signalling linked genes and receptor-encoding genes contributed to the donor-specific variability (PC3 loadings, **Suppl. Table 4**). Individual PC loadings values were moderate (∼0.18 – 0.26, **Suppl. Table 4**), indicating that variance was distributed across multiple genes.

Unstimulated SF and SF treated with TNF + sIL-6 exhibited high inter-sample similarity in gene expression profiles. By contrast transcriptional responses were considerably more variable in IL-6 + sIL-6R–stimulated cells in line with PC3 loading results (**Fig. 5e, Suppl. Table 4**). We identified two major gene expression clusters, generally separating downregulated and upregulated genes in TNF + sIL-6R–stimulated SF (**Fig. 5f**). Downregulated genes (Gene Module 0) included *STAT5B*, SMAD family genes as well as genes, encoding transferrin receptor, JAK2–STAT5-linked receptors (37) and JAK–STAT pathway regulators (**Fig. 5f, Suppl. Fig. 2c**). In the upregulated gene cluster, three distinct gene modules (Modules 1-3) emerged (**Fig. 5f**). Gene module 1 was strongly induced in TNF + sIL-6R–stimulated SF. The Module 1 comprised inflammatory genes (*MMP3*), genes encoding the ISG3 complex components (*STAT1, STAT2*, *IRF9*), and ISGF3-dependent canonical interferon-response genes (*GBP1*, *IRF1*, *ISG15*, *OAS1*, **Fig. 5f, Suppl. Fig. 2d**), highlighting robust inflammatory and interferon-driven signaling. Gene modules 2 and 3 contained the core IL-6–linked JAK–STAT pathway genes (*IL6ST*, *JAK2*, *STAT3*, *SOCS3*), alongside genes encoding different transcriptional regulators *CEBPB*, *JUNB*, and *MYC* (**Fig. 5f, Suppl. Fig. 2d**). Genes in the Modules 2 and 3 were upregulated in both TNF + sIL-6R– and IL-6 + sIL-6R–treated SF (**Fig. 5f, Suppl. Fig. 2d**).

Together, these data highlighted stimulus-biased engagement of interferon-versus IL-6–centered JAK–STAT programs in SF.

### IL-6 and TNF drive overlapping and distinct signaling pathways in SF

We further dissected TNF– and IL-6-driven transcriptional responses in SF. Cultured SF were stimulated with IL-6 + sIL-6R, TNF + sIL-6R, or TNF alone. TNF activates multiple pathways, including NF-κB and AP-1. Thus, we analysed the expression of additional genes (*IL8*, *MMP3*, *TNFAIP3*) linked to these pathways. qPCR confirmed both a shared (*IL6ST*, *JAK2*, *STAT3*, *SOCS3*, *IRF9*, *IL-6*, **Fig. 6a**) and stimulus-biased (*JAK3*, *STAT2*, *IRF1*, *ISG15*, *OAS1*, *GBP1*, **Fig. 6b**) gene expression of the core JAK–STAT pathway components and interferon response genes. Increase in *SOCS3* and *JUNB* expression was strongest in the presence of sIL-6R (**Fig. 6a**). JAK3, minimally expressed in unstimulated SF, was significantly induced by TNF (**Fig. 6b**). This result suggested a possible TNF-driven stromal JAK3 expression in TNF-rich microenvironments. Among pro-inflammatory, regulatory, and matrix-remodeling genes, *IL-6* and *MMP1* were upregulated by IL-6 + sIL-6R and TNF, with or without sIL-6R (**Fig. 6a**). By contrast, *NFKB1*, *TNFAIP3*, *IL8*, and *MMP3* genes were induced by TNF (**Fig. 6b**) independently of sIL-6R.

In summary, SF activated both overlapping and distinct signalling pathways and gene expression programs in response to different inflammatory stimuli.

### SF show potent synergy between IL-6 trans-signaling and TNF pathways

Various proinflammatory stimuli can enhance fibroblast secretion of IL-6 and IL-8, the key mediators in RA pathogenesis. SF stimulated with 0.1ng/ml TNF + sIL-6R secreted IL-6 at levels comparable to 1ng/ml TNF alone, with further enhancement at 1ng/ml TNF + sIL-6R (**Fig. 6c**). In contrast, IL-8 secretion was solely dependent on TNF concentration, with no synergistic effect of sIL-6R (**Fig. 6d**).

### Cytokine context shapes tofacitinib effects on STAT1/3 phosphorylation

Tofacitinib is a reversible pan-JAKi that primarily inhibits JAK1/3 kinases and to a lesser extent JAK2. In *in vitro SF* studies tofacitinib is commonly used in concentrations exceeding the therapeutic window in RA patients [reviewed in (16)]. In our experiments, we employed therapeutically relevant tofacitinib concentrations (80nM, 180nM). These concentrations approximate the mean and maximal plasma concentrations achieved with clinically approved dosing regimens in RA (https://www.accessdata.fda.gov, 38-39). The effects were compared to 1000nM tofacitinib, widely used in *in vitro* SF studies.

In IL-6 + sIL-6R-stimulated SF, tofacitinib dose-dependently suppressed STAT1 and STAT3 phosphorylation (**Fig. 7a-d**). At 180nM concentration, tofacitinib reduced pSTAT1 and pSTAT3 levels by 98.5% and 80.9%, respectively, at 10min time point (**Fig. 7e**), and by 89.6% and 66.4%, respectively, at 5h (**Fig. 7f**) of IL-6 + sIL-6R stimulation. Suppression was strongest for STAT1 phosphorylation, and the inhibitory effect on pSTATs declined over time. A significant, albeit small, decrease in pJAK1 was observed in SF treated with supraphysiological 1000nM tofacitinib at 10min of IL-6 + sIL-6R. No decrease in pJAK1 was observed at 5h stimulation nor with 180nM tofacitinib. Overall, high JAK1 phosphorylation persisted across both time points in IL-6 + sIL-6R-stimulated SF (**Fig. 7a-d**).

In SF stimulated with TNF + sIL-6R and TNF alone, 180nM tofacitinib significantly inhibited cytokine-induced pSTAT1 (80.4% and 87.7%, respectively, **Fig. 7b, d, f**) and nearly maximally suppressed pSTAT3 (94.1% and 94.7%) (**Fig. 7b, d, f**). These results contrasted with IL-6 + sIL-6R stimulation. They likely reflected smaller amounts of autocrinally-produced IL-6 driving pSTAT3 (**Fig. 7b**), alongside increased basal STAT1 phosphorylation at 5h.

### Persistent STAT1/3 Phosphorylation Under Cytokine Load Suggests a JAK-Inhibitor Ceiling

Despite efficient suppression of STAT phosphorylation, residual pSTAT1 and pSTAT3 persisted in cytokine-stimulated tofacitinib-treated SF, particularly at the therapeutic tofacitinib concentration (**Fig. 7g, h**). This residual phosphorylation was stimulus-dependent; at 180nM tofacitinib concentration, pSTAT1 was detected across all cytokine conditions, while pSTAT3 was prominent only in IL-6 + sIL-6R-stimulated SF (**Fig. 7a, g, h**).

### Tofacitinib spares *TNFAIP3* expression while suppressing SOCS3 and proinflammatory gene expression

We next assessed tofacitinib’s effects on selected JAK–STAT-pathway associated genes (*SOCS3, ISG15*), TNF-signalling-linked genes (*TNFAIP3*), and fibroblast function-linked genes (*IL6*, *IL8*, *MMP1*, *MMP3*). Tofacitinib suppressed IL6, MMP1 and MMP3 mRNA expression across cytokine stimuli. These genes were strongly induced by TNF and their suppression by tofacitinib, albeit significant, remained partial. ISG15 mRNA levels decreased in TNF + sIL-6R-stimulated, tofacitinib-treated SF (**Fig. 8a**). The expression of *SOCS3* was decreased by tofacitinib in IL-6/TNF + sIL-6R-treated SF (**Fig. 8a**). IL8 and TNFAIP3 transcript levels were unaffected by tofacitinib treatment (**Fig. 8a**), indicating that these genes are not primary downstream targets of the JAK–STAT pathway in SF.

**Figure 8.**
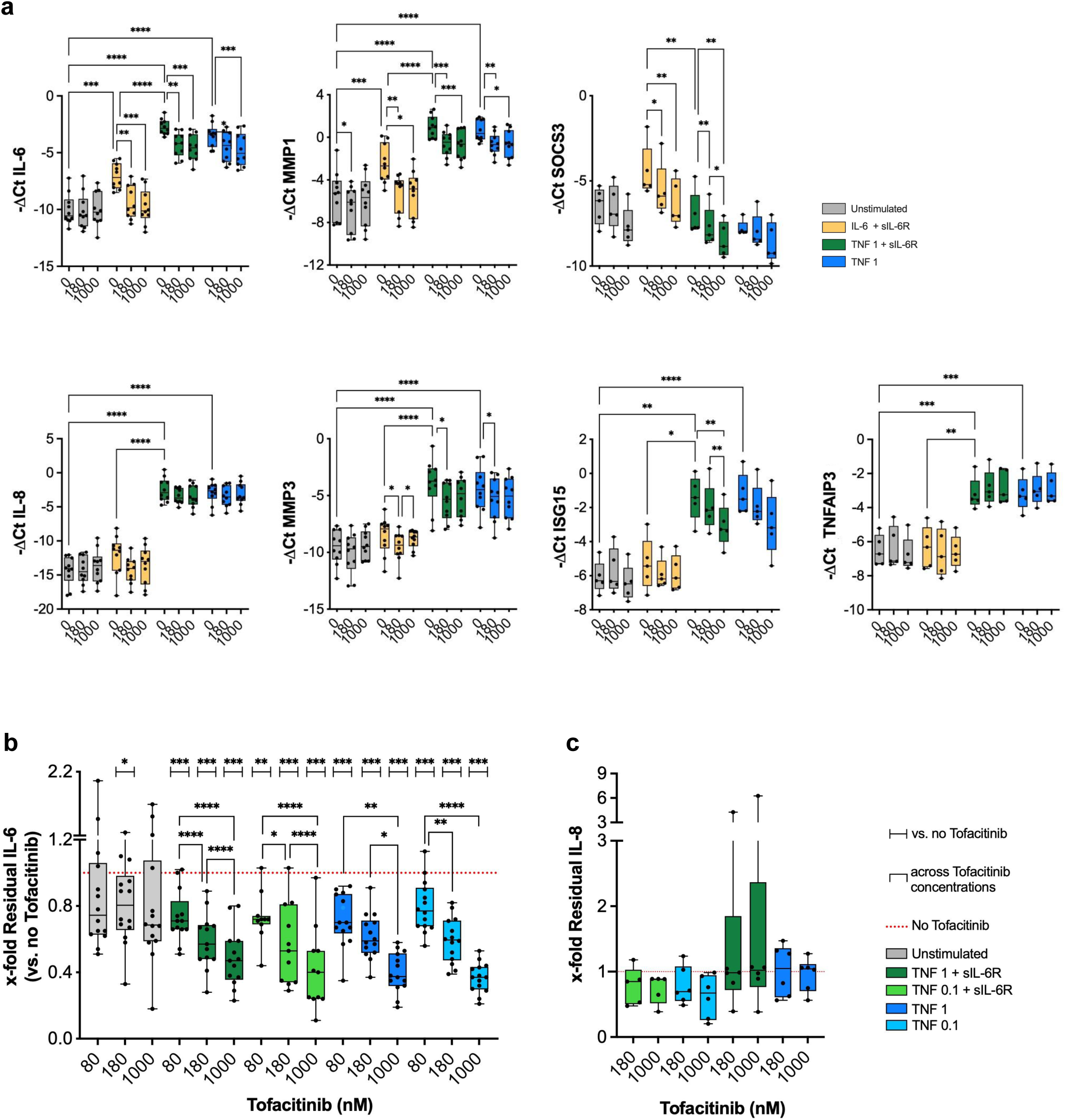
Cytokine microenvironments shape tofacitinib effects on inflammation in rheumatoid arthritis synovial fibroblasts. **a)** *In vitro* experiments. We stimulated cultured rheumatoid arthritis (RA) synovial fibroblasts (SF) with 50ng/ml interleukin 6 (IL-6) or 1ng/ml tumor necrosis factor (TNF 1) in the presence or absence of 50ng/ml soluble IL-6 receptor (sIL-6R) for 24h across tofacitinib concentrations (0, 180, 1000nM). Cells were pretreated with tofacitinib for 2h. Gene expression was quantified using qPCR and calculated as ΔCt. N = 5-10 biological replicates per experimental condition. Data are presented using box and whiskers plots with median and min to max. To compare unstimulated and stimulated experimental conditions in the absence of tofacitinib, we performed RM one-way ANOVA with Šídák’s multiple-comparisons test or the Friedman test with Dunn’s multiple-comparisons test. Biologically meaningful pairwise comparisons were performed. To compare tofacitinib concentration effects within individual experimental conditions (unstimulated and stimulated), we used RM one-way ANOVA with Tukey’s multiple-comparisons test or the Friedman test with Dunn’s multiple-comparisons test. **b, c)** *In vitro* experiments. We stimulated cultured RA SF with 50ng/ml IL-6, 0.1 or 1ng/ml TNF in the presence or absence of 50ng/ml sIL-6R across tofacitinib concentrations (0, 80, 180, 1000nM). Cells were pretreated with tofacitinib for 2h. Secreted **b)** IL-6 (n=11-14 biological replicates) and **c)** IL-8 (n=5-6 biological replicates) proteins were analysed 24h after stimulation using ELISA. Data are presented using box and whiskers plots with median and min to max. *x-fold = Interleukin(s,d) / Interleukin(s,0)*, where *Interleukin* denotes the cytokine (IL-6, IL-8), *s* denotes the experimental condition (unstimulated or stimulated), *d* denotes the tofacitinib concentration (0, 80, 180, 1000nM), and x-fold = 1 represents the baseline x-fold, calculated for the 0nM tofacitinib for each experimental condition. To compare x-fold values to baseline x-fold = 1, we performed a one-sample t-test or Wilcoxon test. p-values were adjusted for multiple comparisons with the Bonferroni test. To compare x-fold values across tofacitinib concentrations within individual experimental conditions, we used RM one-way ANOVA with Tukey’s multiple comparisons test or Friedman test with Dunn’s multiple comparisons test. *p<0.05, **p<0.01, *** p<0.001, ****p<0.0001.

### IL-6 secretion persists in tofacitinib-treated TNF-stimulated synovial fibroblasts

To further elucidate tofacitinib’s divergent effects on proinflammatory cytokines in SF, we quantified IL-6 and IL-8 secretion. SF were stimulated with 0.1ng/ml or 1ng/ml TNF ± sIL-6R, mimicking distinct cytokine quantities and synergies in the RA joint milieu.

Tofacitinib dose-dependently inhibited IL-6 secretion by ∼20–30%, 50% and 60% at 80, 180 and 1000nM concentrations, respectively, across stimuli (**Fig. 8b**). At therapeutically relevant concentrations (80nM, 180nM), tofacitinib-driven IL-6 inhibition was partial (**Fig. 8b**). Absolute IL-6 secretion varied with stimulus quantity and synergy (**Fig. 6c**), whereas percent inhibition of IL-6 secretion depended solely on tofacitinib concentration (**Fig. 8b**). Consequently, higher residual IL-6 was detected in SF under stronger TNF stimulation or sIL-6R co-exposure (**Fig. 8b**). These data further underscored potential ceiling effect of therapeutic tofacitinib concentrations in severely inflamed RA joints.

Consistent with prior reports (33) and gene expression data (**Fig. 8a**), IL-8 secretion remained largely insensitive to tofacitinib (**Fig. 8c**), suggesting regulation by tofacitinib-independent pathways (40).

### Acute tofacitinib withdrawal results in STAT1/3 re-phosphorylation in rheumatoid arthritis synovial fibroblasts

In IL-6 + sIL-6R–stimulated SF, tofacitinib did not diminish JAK1 phosphorylation, which remained high at both 10min and 5h of stimulation (**Fig. 7a-d**). This prompted us to examine whether pJAK1 could acutely re-activate STAT1/3 when tofacitinib and stimuli are withdrawn. Accordingly, we washed out tofacitinib and stimuli after 10min or 5h of stimulation, followed by pJAK1 and pSTAT1/3 detection at 10min after the washout. Experimental design is shown on **Fig. 9a**.

**Figure 9.**
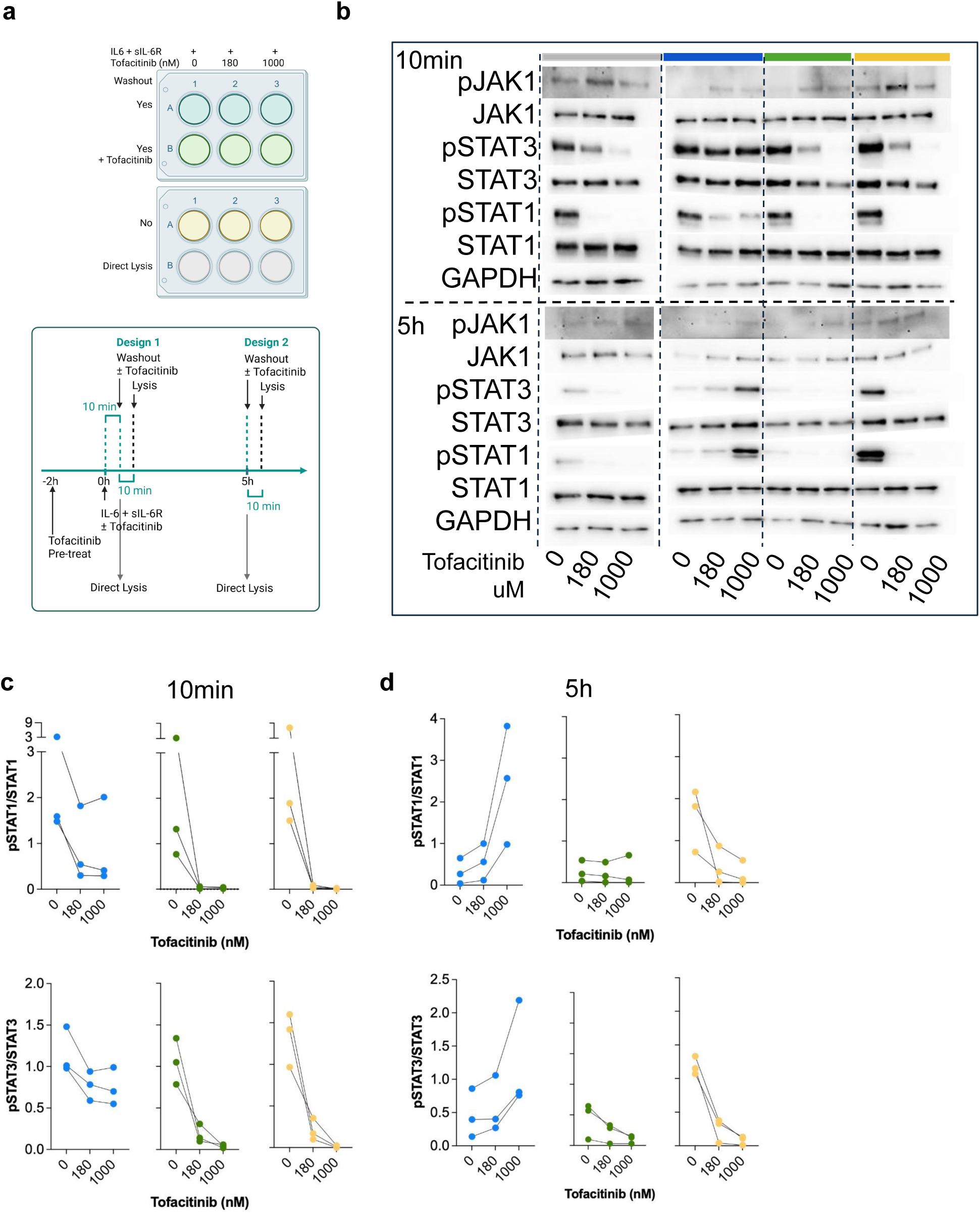
STAT phosphorylation rapidly resurges in rheumatoid arthritis synovial fibroblasts following acute tofacitinib withdrawal. **a)** Schematic of the tofacitinib washout experiment in cultured rheumatoid arthritis synovial fibroblasts (SF); created with BioRender. We pretreated SF with tofacitinib (180, 1000nM) or equal volumes of DMSO for 2h. Thereafter, we stimulated SF with 50ng/ml interleukin 6 (IL-6) and 50ng/ml soluble IL-6 receptor (sIL-6R) in the presence of tofacitinib (180, 1000nM) or DMSO for 10min or 5h. Finally, we either directly lysed SF, washed out supernatants with/without tofacitinib re-supplementation, or continued treatment for an additional 10min. **b)** Representative immunoblot images of (un)phosphorylated JAK1 and STAT1/3 proteins and GAPDH (loading control) in SF are shown across experimental conditions. Original immunoblot images are depicted in **Suppl. Figs. 5, 6**. We quantified experimental data from **c)** 10min and **d)** 5h washout experiments as a ratio of phosphorylated versus total protein, using optical density values of chemiluminesce-detected immunoblotted proteins. N=3 biological replicates per experimental condition.

Drug-stimuli washout triggered a rapid pSTAT1 and pSTAT3 rebound (**Fig. 9b-d**). pSTAT1/3 rebound patterns, however, differed between the 10min and 5h time points. In 10min–stimulated SF, pSTAT1/3 rebound was evident in tofacitinib-treated cells, but strongest signals were detected in tofacitinib-naive cells (**Fig. 9b, c**). These findings suggested that IL-6 and sIL-6R remained membrane-bound for a longer time period whereas intracellular tofacitinib concentrations declined rapidly upon drug removal. By contrast, at 5h timepoint STAT1/3 re-phosphorylation escalated with tofacitinib dose (**Fig. 9b, d**). Re-supplementing tofacitinib efficiently rescued the pSTAT1/3 rebound at both time points (**Fig. 9b-d**).

JAK1 phosphorylation was detected across experimental conditions and time points in the washout experiments (**Fig. 9b, Suppl. Figs. 5, 6**). However, pJAK1 bands were faint in the washout experiments (**Fig. 9b; Suppl. Fig. 6** compared with **Fig. 7a,b**), particularly at 5h, precluding reliable quantification. Reduced pJAK1 signal coincided with lower total JAK1 and GAPDH signals (**Fig. 9b; Suppl. Fig. 6** compared with **Fig. 7a, b**), consistent with smaller cell input in the washout experiments.

## Discussion

Synovial fibroblasts – central drivers of chronic inflammation in RA – are increasingly linked to D2T RA (3, 12, 13, 28, 30). The lack of therapies that directly target fibroblasts may partly explain this association, as most RA treatments affect SF only indirectly via immune cell modulation. JAKi are a notable exception, acting on both structural and immune cell-driven pathways.

Here we provide substantial evidence for clinical observations that JAK inhibitors – compared to other DMARDs (3, 28, 30) – achieve greater reduction in arthritis activity in patients with D2T RA (14, 41). Our findings highlight broad JAK1 expression across synovial pathotypes, cell populations, and SF subsets. Notably, JAK1 is the dominant Janus kinase in all SF subsets, including DKK3□ and CD34□ SF enriched in drug-refractory RA. Therefore, JAK1-selective and pan-JAK inhibitors may directly target fibroid synovial tissue as well as DKK3□ and CD34□ SF in D2T RA. *In vitro* experiments with the pan-JAK inhibitor tofacitinib corroborated the effective SF targeting, further implying that SF are direct pharmacological targets of JAK inhibition in RA.

Our results align with growing synovial biopsy–driven, precision medicine efforts to stratify RA patients into therapeutic subgroups based on responsiveness to distinct treatment modalities. Specifically, transcriptomic analyses of JAK-STAT signalling networks inferred that CXCL12^high^, HLA-DR+ and PRG4+ SF subsets are likely to be particularly JAKi sensitive. CXCL12^high^ and HLA-DR+ SF displayed the broadest and strongest JAK-STAT pathway-linked gene expression, featuring the activation of both TNF/type I interferon–driven and especially IL-6–driven signaling. The IL-6-driven JAK-STAT gene program appeared autocrinally potentiated through specific expression of IL-6 in these cells, which was further validated in our *in vitro* experiments. Aligned with these findings, CXCL12^high^ and HLA-DR^high^ SF were enriched in T cell – fibroblast-CTAP as well as myeloid cell-dominated CTAP – high in *TNF* and interferon gene expression (11). Upadacitinib blocked IFNγ-driven HLA-DR induction in SF (42). Consistent with SF transcriptomic signatures, SELE^+/high^ synovial venous endothelial cells were enriched in IL-6 and SOCS3 transcripts (4), supporting an expanded model of autocrine IL-6 signaling within the synovial structural cell compartment.

By contrast, lining PRG4□ SF prominently expressed *STAT1* and interferon response genes but not TNF receptor genes, likely preferring a direct interferon-driven JAK-STAT signalling. Localisation of STAT1 to synovial lining was previously reported (32). Lining SF are key mediators of cartilage and bone destruction in RA (5) and their JAKi sensitivity could mirror the clinical efficacy of JAKi in erosive RA. Systematic analysis of the effects of JAK inhibitors with varying selectivity profiles will be crucial to validate the projected vulnerabilities of SF subsets to JAKi.

We demonstrated that the inflammatory context shapes both JAK–STAT signaling and tofacitinib responses in SF. This is particularly relevant given interpatient variability in synovial cell composition, shaping in turn the joint’s pro-inflammatory milieu and fibroblast activity states (8, 11). pJAK1 was strongly induced in IL-6 + sIL-6R treated SF, but not in TNF ± sIL-6R–stimulated SF. TNF could induce subthreshold pJAK1 levels – not detectable with immunoblotting, resulting from limited autocrine IL-6 production at 5□h. IL-6 dose titration would be necessary to establish the IL-6 threshold required for reliable pJAK1 detection in SF. Measuring pJAK1 at later time points in TNF ± sIL-6R–stimulated SF may provide additional information. Alternatively, TNF may preferentially utilise alternative JAKs in SF, notably JAK2 (43).

Our study uncovered strong synergy between soluble IL-6 receptors and TNF in SF. The TNF–IL-6 co-signaling was shown to amplify inflammatory responses in co-cultured SF and T cells (44). A bispecific anti-TNF/IL-6 nanobody disrupted the SF-T cell circuits, maintaining remission in a collagen-induced arthritis (44). Additionally, TNF cooperated with oncostatin M in SF (45). This synergy could provide a mechanistic explanation for why JAKi and IL-6R blockade often outperform TNF inhibitors in patients with refractory RA (14, 41). TNF inhibitors may not fully break the sIL-6R-driven amplification loop in SF, whereas JAKi and IL□6R blockade potentially could. D2T-RA resolved more frequently in patients treated with tocilizumab (41), which targets both soluble and membrane-bound IL-6R. Collectively, these data underscore the role of cytokine cooperation in RA pathogenesis and refractory RA. Targeting multiple cytokines or cell types could help overcome treatment resistance. This aligns with the evidence for an enhanced efficacy of JAKi (14) and bi-specific T cell engagers (46) in D2T RA.

Our data unraveled a functional JAKi ceiling effect for STAT1/3 phosphorylation and IL-6 secretion in SF within cytokine-rich or –synergistic milieus. Specifically, residual IL-6 and pSTAT1/3 activities were detected in highly inflamed SF at therapeutic tofacitinib doses, likely reflecting a dose trade-off between drug efficacy and safety. Cytokine concentrations in our *in vitro* experiments were kept within pathologic ranges observed in RA patients, mimicking highly inflamed joints. We approximated mean sIL-6R concentration in RA synovial fluid (36) and mimicked synovial fluid TNF levels in moderate and active/severe RA (47, 48). Experimental IL-6 concentrations captured the upper synovial fluid concentration ranges in RA (49). The functional JAKi ceiling effect inferred that in patients with high baseline cytokine activity, achieving deeper SF suppression might require higher JAKi exposures. Higher JAKi doses are constrained in clinical practice by safety considerations (18,19). Moreover, suppression of SF activation in RA joints would depend not only on baseline cytokine activity – baseline inflammation and tofacitinib dose, but also on the drug’s ability to inhibit other cytokine-producing synovial cell types (50). Clinically, inflammation burden was reduced with duration of tofacitinib treatment (51, 52), which could mirror, at least partially, the drug’s cumulative effect on multiple cell types.

Synovial pSTAT decreased in tofacitinib treated RA patients (53); pSTAT blockade, however, remained partial and stimulus-/cell-type dependent *in vivo*, even after months of therapy (54). High baseline disease activity was identified as a predictive factor for developing D2T-RA (3). Moreover, pan-JAKi-treated patients with high baseline disease activity had significantly lower JAKi retention rates (55). Therefore, the functional tofacitinib ceiling effect offers a plausible mechanistic basis for a partial therapeutic response and lower tofacitinib compliance in RA patients with high baseline inflammation despite adequate dosing. Both, the ceiling effect as well as the benefit/risk trade-off could contribute to tofacitinib’s lower retention rates in RA patients with high baseline arthritis activity.

Our study revealed an intrinsic decoupling mechanism between pJAK1 and pSTAT1/3 in inflamed SF. Whereas JAK1 phosphorylation remained largely intact in IL-6 + sIL-6R–stimulated SF, STAT1/3 phosphorylation declined over experimental time. These findings suggested an activation of counter-regulatory mechanisms mediated by negative pathway regulators (31, 32). Indeed, inflamed SF strongly upregulated SOCS3, potentially serving as an intrinsic brake on pro-inflammatory amplification. Analogously, pJAK-pSTAT1/3 uncoupling was evident in inflamed SF treated with therapeutic tofacitinib doses. Specifically, 180nM tofacitinib suppressed STAT1/3 phosphorylation without pJAK1 suppression. Similarly, stable JAK2 phosphorylation accompanied by effective pSTAT inhibition was reported in cells harboring activating JAK2 mutations or exhibiting cytokine-induced JAK2 activation (56). By contrast, 300nM tofacitinib partially reduced JAK1 phosphorylation in oncostatin M-stimulated cells. In our experiments, superphysiological 1000nM tofacitinib caused a small, albeit significant pJAK1 reduction at 10min of IL-6 + sIL-6 stimulation. These data implied that drug dose and stimulus identity may influence pJAK1 sensitivity to tofacitinib (57, 58). Analysis of diverse stimuli, other Janus kinases and differentially selective JAK inhibitors could further clarify JAK-STAT vulnerabilities in SF.

The decoupling between persistent pJAK1 and declining pSTAT can have critical implications in JAKi withdrawal phenomenon and JAKi tapering in a clinical setting. It suggests a “primed kinase, restrained transcription” state, causing a rapid pSTAT resurgence upon abrupt JAKi withdrawal. Specifically, SF stimulated with IL-6 + sIL-6R displayed an acute pSTAT rebound upon tofacitinib washout. pSTAT rebound was particularly strong after prolonged tofacitinib exposure and at high tofacitinib concentrations. Tofacitinib re-administration efficiently rescued pSTAT1/3 rebound, consistent with suppression of withdrawal-linked RA symptoms upon tofacitinib re-initiation (20). In mononuclear cells from myelofibrosis patients, ruxolitinib facilitated phosphorylation of the activation loop, stabilizing JAK2 in its active kinase conformation (59). Additionally, ruxolitinib inhibited pJAK2 degradation, further enhancing pSTAT1/3/5 signaling upon drug withdrawal (59). Type I JAKi, such as tofacitinib and ruxolitinib, can bind and stabilize an active JAK conformation, underlying JAK inhibitor withdrawal syndrome (56, 59). Whether this mechanism operates along the JAK1–STAT1/3 axis in SF remains to be determined. In our washout experiments, pJAK1 bands were faintly detected by immunoblotting, precluding robust quantification. As a result, the hypothesis that pJAK1 becomes increasingly locked in an active conformation with prolonged tofacitinib exposure could not be rigorously tested.

Rebound effects were monitored in SF for a short period post-tofacitinib cessation; such a granular timeline is challenging to capture in a real-life clinical scenario. Nonetheless, abrupt tofacitinib withdrawal in RA patients was associated with acute arthritis worsening, increased adverse events, and higher flare incidence (20, 60). Similarly, ruxolitinib discontinuation in myelofibrosis patients triggered a rare but serious ruxolitinib withdrawal syndrome, with flaring myelofibrosis symptoms and signs (61). Flares emerged from 24h to three weeks post drug discontinuation, attributed to resurgent JAK2–STAT signaling (62, 63). Thus, ruxolitinib should be preferably discontinued in a tapering schedule and under close physician supervision (61). A gradual tapering of JAKi is recommended also in patients with RA after achieving a remission or low disease activity, with reintroduction of the last effective JAKi dose upon RA flare (64).

Strikingly, tofacitinib repressed the IL-6–driven *SOCS3* gene expression. This reduction may weaken intrinsic SOCS3-mediated negative feedback, thereby contributing to sustained JAK1 phosphorylation (65) and pSTAT rebound in inflamed SF. Consistent with our findings, studies in PBMCs from RA patients showed that tofacitinib decreases SOCS3 mRNA expression (54). Based on these observations, we hypothesized that preventing context– or time-dependent SOCS3 suppression may help avert pSTAT rebound and enhance the drug’s direct anti-inflammatory efficacy. However, because SOCS3 protein levels have been reported to increase in tofacitinib-treated SF (66), measuring SOCS3 protein across relevant contexts would be necessary to substantiate our hypothesis.

Whereas *SOCS3* gene expression decreased in the presence of tofacitinib, *TNFAIP3* expression was not affected.This suggests that in tofacitinib-treated SF, the capacity to negatively regulate TNF signaling (67) is preserved. A hypothesis that merits further study, particularly considering the safety profile of JAKi versus other biological DMARDs.

Finally, the washout experiments suggested a rapid loss of tofacitinib activity in SF upon acute drug removal, likely reflecting fast intracellular efflux of tofacitinib from these cells (68).

In summary, this study highlights broad expression of JAK1 across synovial cell types, SF subsets and synovial pathotypes, underscoring the potential for JAK1 selective and pan-JAK inhibitors to address the interpatient synovial cell/fibroblast heterogeneity in RA. We emphasise potent synergy between TNF and IL-6 transsignaling, escalating fibroblast IL-6 secretion; and underscore an adaptive uncoupling of STAT1/3 from persistent JAK1 activation that may restrain fibroblast-driven inflammatory activities. Under high cytokine conditions, residual STAT1/3 phosphorylation and IL-6 secretion persists, suggesting a functional ceiling effect of pan-JAK inhibitor tofacitinib in SF, and abrupt drug withdrawal triggers rapid STAT1/3 reactivation. Collectively, our results position SF as key cellular targets of JAK inhibition in RA and resolve JAK–STAT pathway dynamics in these cells, informing both therapeutic benefit, limitations and withdrawal-related risk of tofacitinib therapy in RA.

## Acknowledgement

We would like to thank the patients for donating synovial tissue. For technical support, we thank Benvinda Henriques Campos and Peter Kuenzler; for critical manuscript reading we thank Tim Killen, Janine Lueckgen and Katja Lakota. This study was funded by the AbbVie Rheumatology Grant, the OPO-Foundation, the Novartis Foundation for medical-biological research and the Vontobel Foundation awarded to MFB. BB was supported by the Agency of Research and Innovation Slovenia, National Research Programme “Systemic autoimmune diseases P3-0314. MPB was supported by a scholarship from the Digital Society Initiative (DSI) of the University of Zurich. AŽ, NI and TK were supported by the EULAR Scientific Training Grants. TK was supported by the IUBMB Wood-Whelan Research Fellowship and the Swiss-European Mobility Programme Scholarship.

## Author contributions

MFB conceived and supervised the study. Experiments were performed by AŽ, SGE, BB, TK, NI, RSZ, LMMP and MH. RG and MPB conducted scRNA-seq analyses. MDR supervised scRNA-seq analyses. OD, SSS, CO and MFB organised the study and collected clinical samples and data. BB prepared Western blot imaging data. MPB, AŽ, BB, TK and MFB analysed the data. MFB, MPB, RG and BB interpreted the data. MFB wrote the manuscript with contributions from MPB, AŽ, BB, TK, RG and MDR. All authors contributed to critical discussion, final manuscript drafting and approved the final manuscript.

## Declaration of interests

MFB had received research funding from AbbVie (AbbVie Rheumatology Grant) and Novartis (Novartis Foundation for medical-biological research) and was an employee of BioMed X (Aug 2021 – July 2025) with research funding from Johnson & Johnson Innovative Medicine. OD has/had consultancy relationships with and/or has received research funding from and/or has served as a speaker for the following companies in the last three calendar years: 4P-Pharma, Abbvie, Acepodia, Aera, Amgen, AnaMar, Anaveon, Argenx, AstraZeneca, Avalyn, Boehringer Ingelheim, BMS, Calluna, Cantargia, CSL Behring, EMD Serono, Galderma, Galapagos, Gossamer, Hemetron, Innovaderm, Kali, Lilly, Mediar, MSD Merck, Nkarta, Novartis, Oorja Bio, Orion, Pliant, Prometheus, Quell, Scleroderma Research Foundation, Skyhawk, Tandem, Topadur, UCB and Umlaut.bio. Patent issued “mir-29 for the treatment of systemic sclerosis” (US8247389, EP2331143). Co-founder of CITUS AG. The other authors declare no conflict of interest.

## Declaration of generative AI and AI-assisted technologies in the writing process

During the preparation of this work the author(s) used ChatGPT in order to improve the readability and language of the manuscript. After using this tool/service, the author(s) reviewed and edited the content as needed and take(s) full responsibility for the content of the published article.

## Supplementary Tables

**Supplementary Table 1 Metadata from subjects donating synovial tissue** Provided are demographic, clinical and therapeutic data for patients with RA who donated synovial tissue specimens for omics analyses and *in vitro* experiments l. Data are shown per dataset and experimental setup.

**Supplementary Table 2 Sequences of SYBR Green and TaqMan primers and probes used in qPCR analysis of gene expression**

**Supplementary Table 3 Specifications of antibodies used for immunoblotting Supplementary Table 4 Loadings on principal components** We analysed the expression of JAK-STAT pathway-associated genes using the TaqMan ArrayCards. SF were left either unstimulated SF or were SF stimulated with IL-6 + sIL-6R or TNF + sIL-6R for 24h. N=11 biological replicates per each of 3 experimental conditions. We quantified gene expression using ΔCt, where ΔCt = mean Ct_target gene – mean Ct_housekeeper gene Ubiquitin C (*UBC*); not detected genes were excluded from analysis. We performed principal component (PC) analysis of ΔCt values for 72 genes across 11 donors (N = 33 conditions) using ClustVis software. Loadings on PC1-3s were extracted to identify genes contributing to variability across the respective PCs.

## Supplementary Figure Legends

**Supplementary Figure 1 Transcriptional signature of the JAK-STAT pathway**-**associated genes across synovial fibroblast subsets and pathotypes in rheumatoid arthritis. a, b)** Analysis of JAK-STAT pathway transcriptional signatures in COL1A1+ rheumatoid arthritis synovial fibroblasts (SF, n=21433) from Dataset 3. For Dataset 3, we extracted RA sample scRNA-seq from our synovial scRNA-seq tissue atlas in inflammatory arthritis (BioStudies Database, ArrayExpress (accession number E-MTAB-11791, (4)). **a)** Violin plots showing the expression of genes, encoding the JAK-STAT pathway regulators across SF subsets. **b)** Contribution of SF subsets to total fibroblast *PTPN1* and *PTPN2* gene expression. The subset-specific contributions were calculated as log10-transformed ratios between gene expression proportion (GEX prop.) and SF subset proportion (SF prop.) per sample. A log10 ratio of 0 indicates comparable expression across cell types, whereas positive and negative values indicate over– and under-expressed genes, respectively, in a given cell type versus all cell types. **c)** Expression analysis of interferon A1 and B1 (*IFNA1*, *IFNB1*) genes across synovial tissue pathotypes in 81 synovial tissues from RNA-Seq Dataset 2 (PEAC Cohort). Normalised expression values, extracted from https://peac.hpc.qmul.ac.uk/ (10).

**Supplementary Figure 2 Expression of the JAK-STAT pathway-associated genes in cultured rheumatoid arthritis synovial fibroblasts a, c-d)** Expression of JAK-STAT pathway-associated genes, probed on TaqMan ArrayCards, in **a, b)** unstimulated rheumatoid arthritis synovial fibroblasts (SF) and **b-d)** SF stimulated with 50ng/ml interleukin 6 (IL-6) + 50ng/ml soluble IL-6 receptor (sIL-6R) or 1ng/ml TNF + 50ng/ml sIL-6R for 24h. N=11 biological replicates per experimental condition. **a, c, d)** Data are presented using box and whiskers plots with median and min to max. **a)** Gene expression in unstimulated SF, quantified as ΔCt. **b)** Principal component (PC) analysis of ΔCt values for 72 expressed genes in SF across three experimental conditions, using ClustVis. PC1 captures the largest variance, dominated by a single outlier sample, while PC2 separates samples according to stimulus. **c)** Genes, significantly downregulated in stimulated SF. **d)** Genes upregulated in cytokine-stimulated SF. **c, d)** Plotted are selected genes from Fig. 5f. Gene expression is quantified as x-fold = 2^−ΔΔCt^. To compare the cytokine-driven and baseline (x-fold = 1) gene expression in SF, we performed one sample t-test or Wilcoxon test, with Bonferroni correction for multiple comparisons (black *). To compare gene expression between stimuli, we used paired t-test or Wilcoxon matched-pairs signed-rank test (green *). *p<0.05, **p<0.01, *** p<0.001, ****p<0.0001.

**Supplementary Figure 3 Original uncropped immunoblot blot images presented in** Figure 6a. Protein bands detected using chemiluminescence (WesternBright ECL HRP substrate) and imaged on a Fusion FX imaging instrument (Vilber). Size marker (kDA) shown.

**Supplementary Figure 4 Original uncropped immunoblot blot images presented in** Figure 6b. Protein bands detected using chemiluminescence (WesternBright ECL HRP substrate) and imaged on a Fusion FX imaging instrument (Vilber). Size marker (kDA) shown.

**Supplementary Figure 5 Original uncropped immunoblot blot images presented in** Figure 8b. Protein bands detected using chemiluminescence (WesternBright ECL HRP substrate) and imaged on a Fusion FX imaging instrument (Vilber). Size marker (kDA) shown.

**Supplementary Figure 6 Original uncropped immunoblot blot images presented in** Figure 8c. Protein bands detected using chemiluminescence (WesternBright ECL HRP substrate) and imaged on a Fusion FX imaging instrument (Vilber). Size marker (kDA) shown.

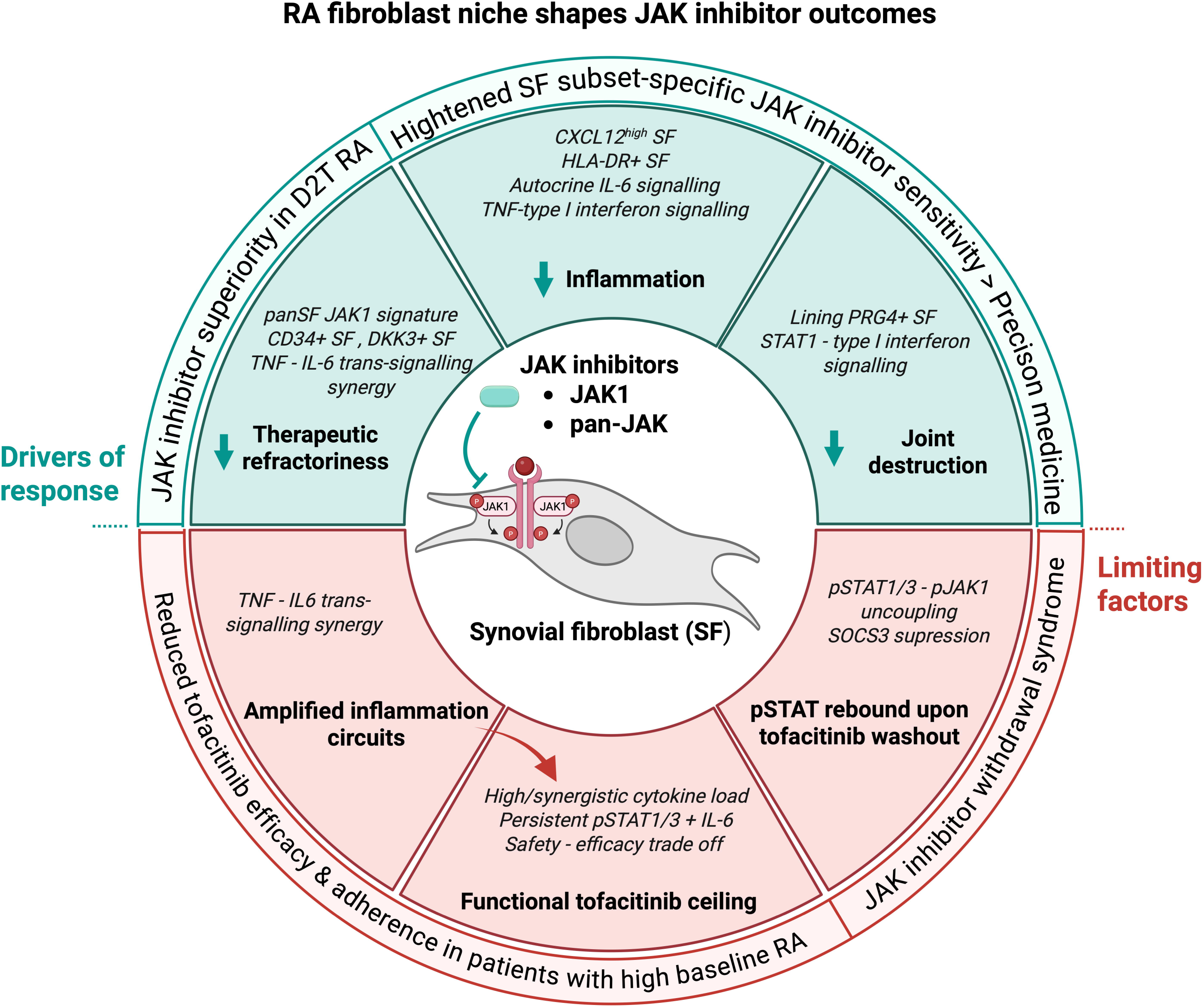

## Notes

https://www.ebi.ac.uk/biostudies/arrayexpress/studies/E-MTAB-11791?query=E-MTAB-11791

https://peac.hpc.qmul.ac.uk

### Competing Interest Statement

Mojca Frank Bertoncelj had received research funding from AbbVie (AbbVie Rheumatology Grant) and Novartis (Novartis Foundation for medical-biological research) and was an employee of BioMed X (Aug 2021 – July 2025) with research funding from Johnson & Johnson Innovative Medicine. Oliver Distler has/had consultancy relationships with and/or has received research funding from and/or has served as a speaker for the following companies in the last three calendar years: 4P-Pharma, Abbvie, Acepodia, Aera, Amgen, AnaMar, Anaveon, Argenx, AstraZeneca, Avalyn, Boehringer Ingelheim, BMS, Calluna, Cantargia, CSL Behring, EMD Serono, Galderma, Galapagos, Gossamer, Hemetron, Innovaderm, Kali, Lilly, Mediar, MSD Merck, Nkarta, Novartis, Oorja Bio, Orion, Pliant, Prometheus, Quell, Scleroderma Research Foundation, Skyhawk, Tandem, Topadur, UCB and Umlaut.bio. Patent issued mir-29 for the treatment of systemic sclerosis (US8247389, EP2331143). Co-founder of CITUS AG. The other authors declare no conflict of interest.

