## Supplemental Figures 1-6 for "Synovial fibroblast niche shapes the efficacy–safety dynamics of JAK inhibition in rheumatoid arthritis": Zupanic et al_Supplemetary Figures_1-6_20260323.pptx

### Slide 1
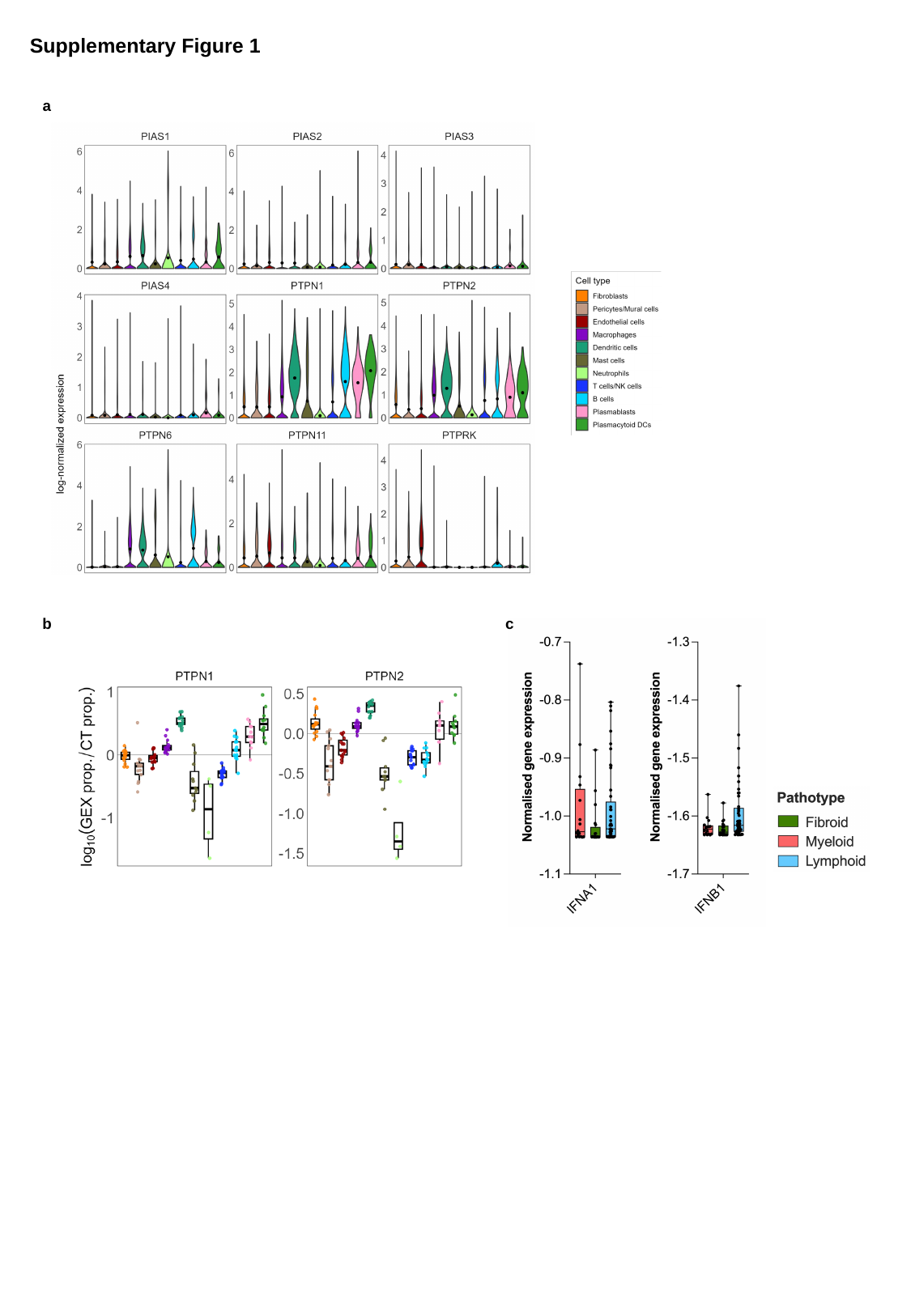

Supplementary Figure 1
a
b
c

### Slide 2
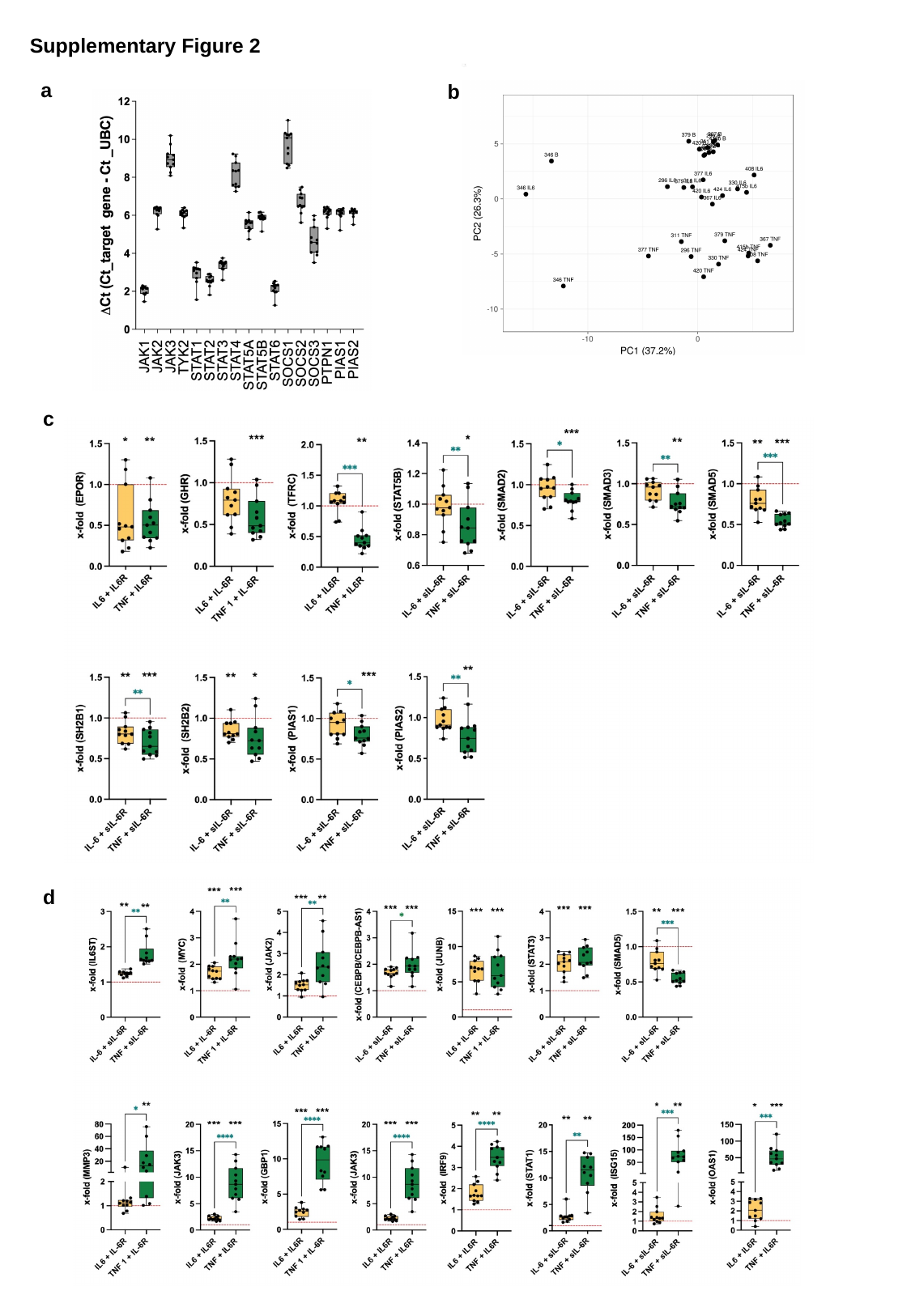

Supplementary Figure 2
a
b
c
d

### Slide 3
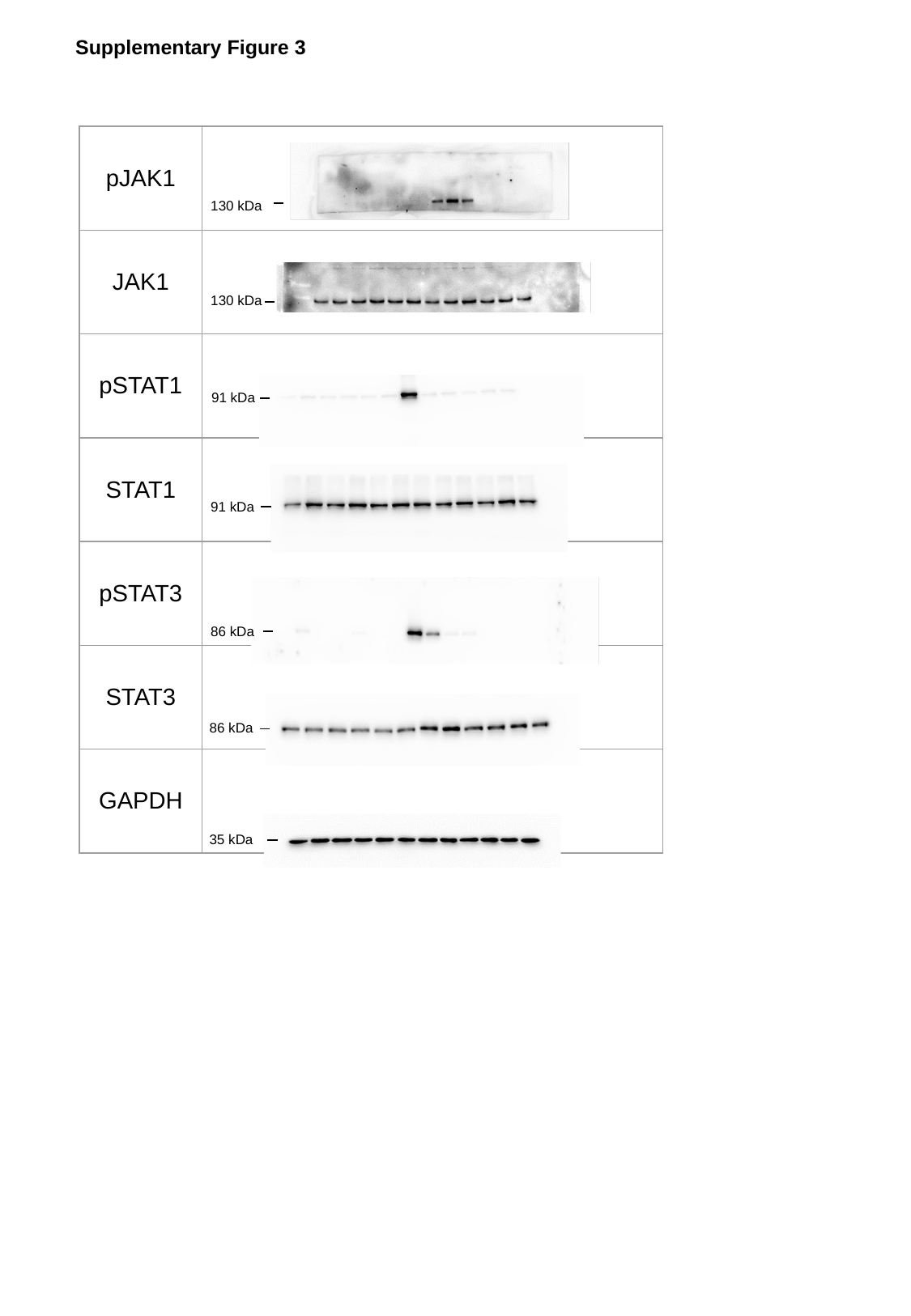

Supplementary Figure 3
| pJAK1 |
| --- |
| JAK1 |
| pSTAT1 |
| STAT1 |
| pSTAT3 |
| STAT3 |
| GAPDH |
130 kDa
130 kDa
91 kDa
91 kDa
86 kDa
86 kDa
35 kDa

### Slide 4
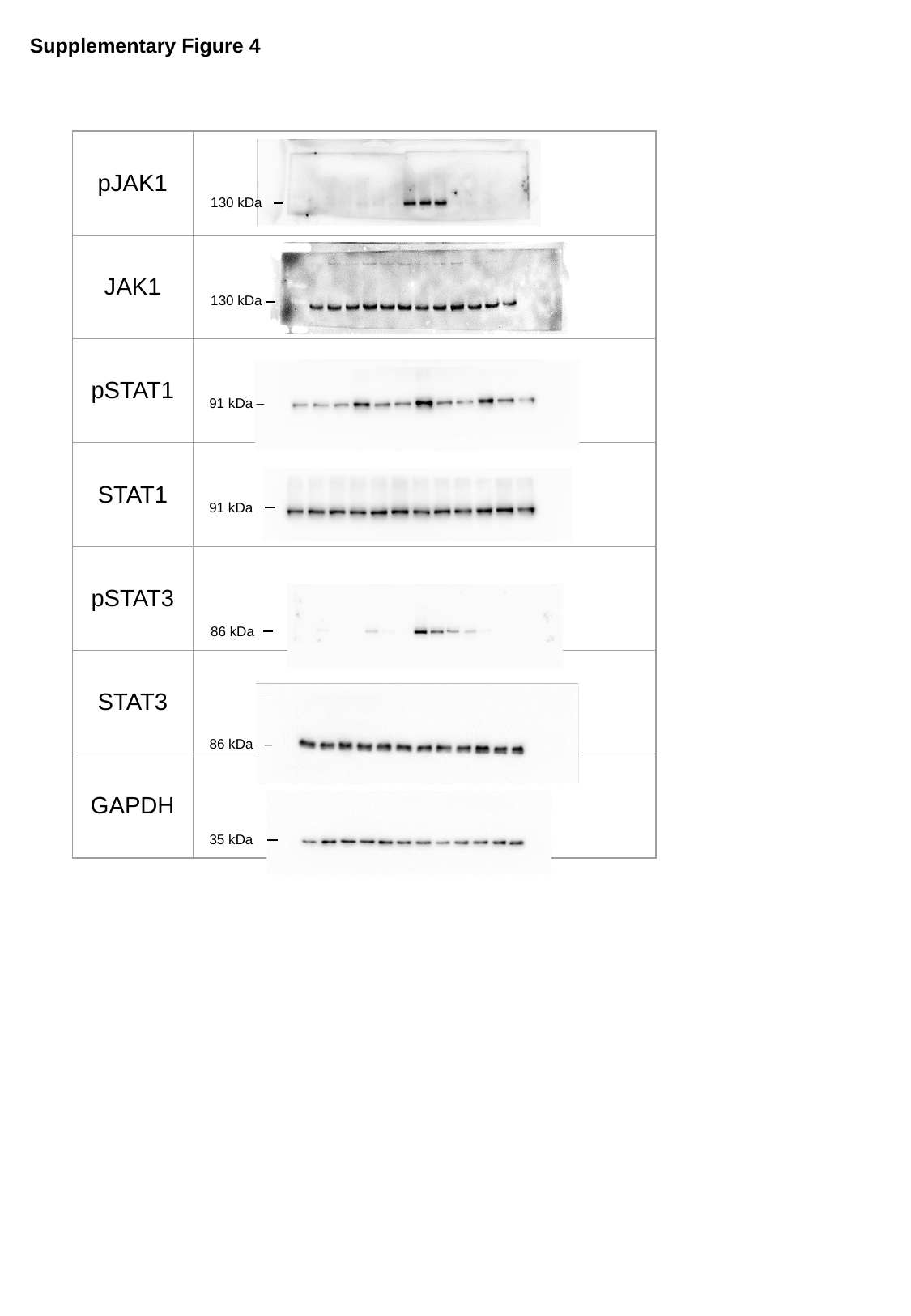

Supplementary Figure 4
| pJAK1 |
| --- |
| JAK1 |
| pSTAT1 |
| STAT1 |
| pSTAT3 |
| STAT3 |
| GAPDH |
130 kDa
130 kDa
91 kDa –
91 kDa
86 kDa
86 kDa –
35 kDa

### Slide 5
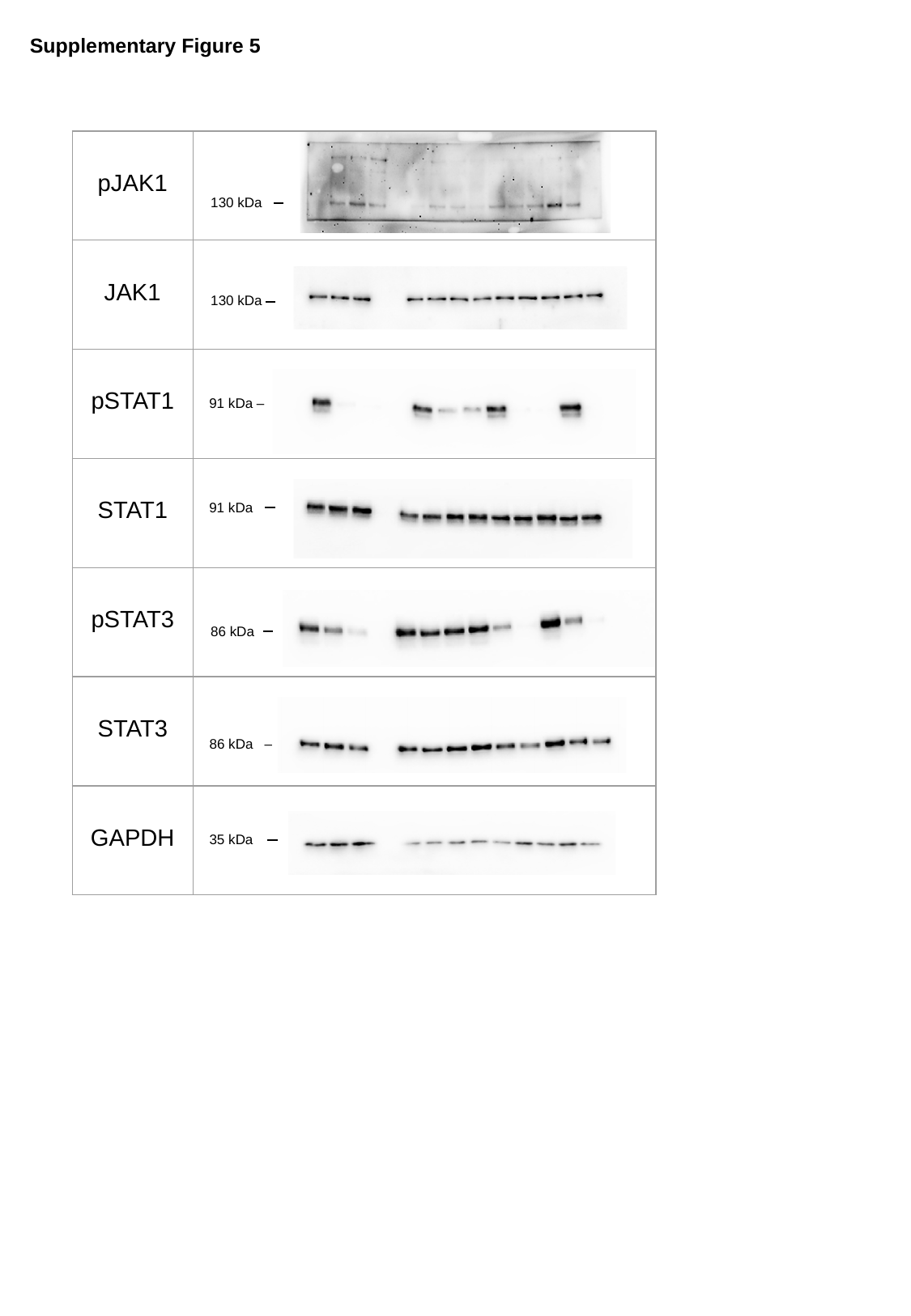

Supplementary Figure 5
| pJAK1 |
| --- |
| JAK1 |
| pSTAT1 |
| STAT1 |
| pSTAT3 |
| STAT3 |
| GAPDH |
130 kDa
130 kDa
91 kDa –
91 kDa
86 kDa
86 kDa –
35 kDa

### Slide 6
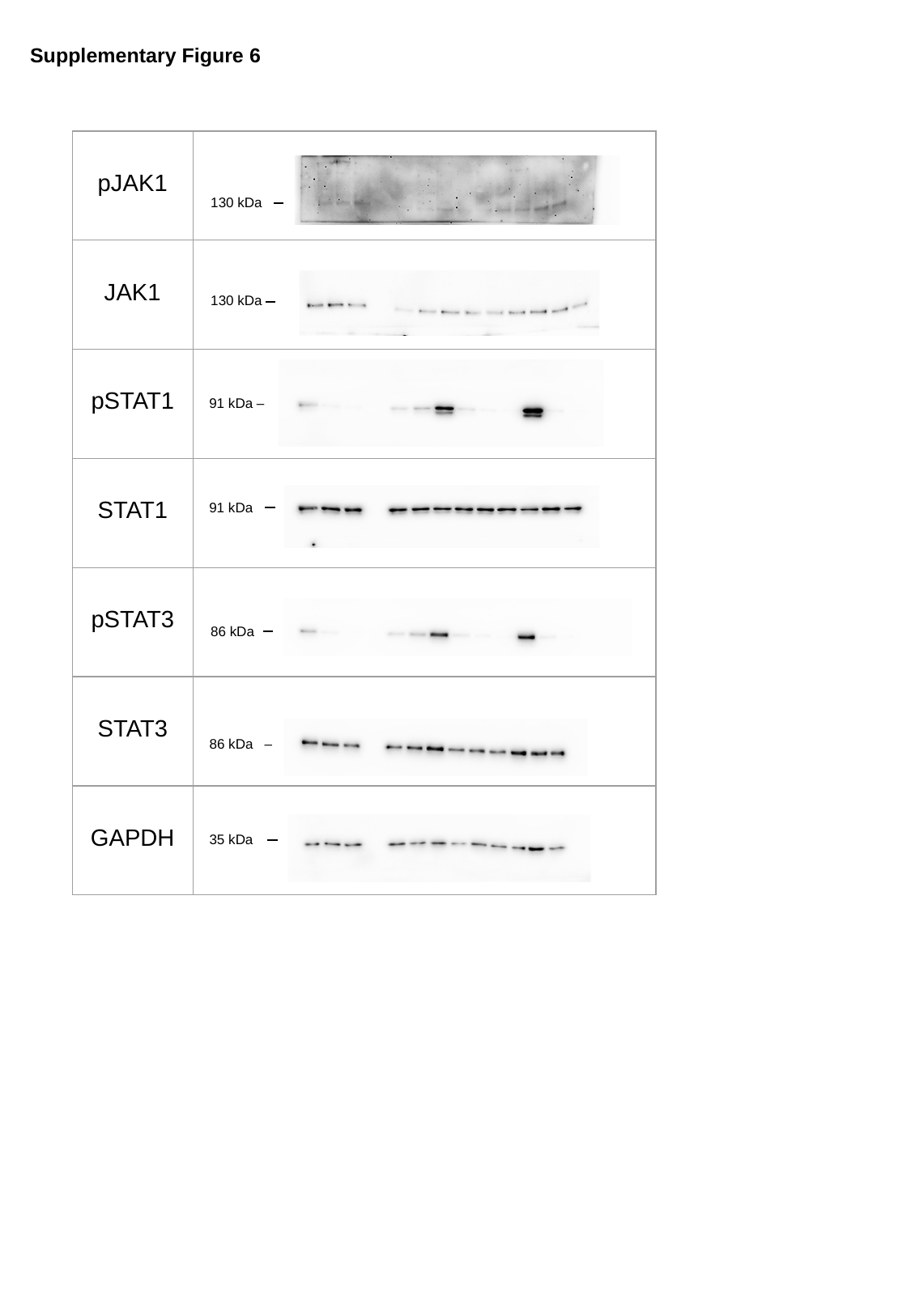

Supplementary Figure 6
| pJAK1 |
| --- |
| JAK1 |
| pSTAT1 |
| STAT1 |
| pSTAT3 |
| STAT3 |
| GAPDH |
130 kDa
130 kDa
91 kDa –
91 kDa
86 kDa
86 kDa –
35 kDa
