## Supplemental Table 3 for "Synovial fibroblast niche shapes the efficacy–safety dynamics of JAK inhibition in rheumatoid arthritis": Zupanic et al_Supplementary Table 3_20260326.docx

**Supplementary Table 3 Specifications of antibodies used for immunoblotting**

| **Target** | **Antibody specification** | **Dilution** |
| --- | --- | --- |
| JAK1 | Rabbit B7125A polyclonal antibody  Assay Biotechnologies | 1:1000 |
| JAK1 pTyr1022 | Rabbit polyclonal antibody A7125  Assay Biotechnologies | 1:1000 |
| STAT1 | Rabbit monoclonal antibody #14995  Cell Signalling Technology | 1:1000 |
| STAT1 pTyr701 | Rabbit monoclonal antibody #9167  Cell Signalling Technology | 1:1000 |
| STAT3 | Mouse monoclonal antibody #9139  Cell Signalling Technology | 1:1000 |
| STAT3 pTyr705 | Rabbit monoclonal antibody #9145  Cell Signalling Technology | 1:1000 |
| GAPDH | Rabbit monoclonal antibody #2118  Cell Signalling Technology | 1:5000 |
