## Supplemental Table 2 for "Synovial fibroblast niche shapes the efficacy–safety dynamics of JAK inhibition in rheumatoid arthritis": Zupanic et al_Supplementary Table 2_20260326.docx

**Supplementary Table 2 SYBR Green Primers**

| **Gene** | **Forward Primer (5' – 3')** | **Reverse Primer (5' – 3')** |
| --- | --- | --- |
| ***STAT2*** | CAGGAAAGGGCAGCTATAAG | TGCAGTTCCTCTGTCACACC |
| ***IRF9*** | AGGAAGGAGGAAGAGGATGC | CAGAGGGACTGAGTGTGCAG |
| ***OAS1*** | TCT GCA CTG TTG CTT TCA GCC AGC | TTG GGA TGG GTC CCC AGT GAG |
| ***NFKB1*** | TCT TAC CCT CAG GTC AAA ATC TG | GAA CAA TAA CCT TTG CTG GTC C |
| ***SOCS3*** | GGG GAA GCA ACA TTT GGA | AAA AGG AGA CCA GCT GAC CA |
| ***JUNB*** | CTC TCA AGC TCG CCT CTT CG | TGC TGT TGG GGA CAA TCA GG |
| ***TNFAIP3*** | AAG GAC AGT GGG CCT GAA ATC | TTC CCC GGT CTC TGT TAA CAA G |
| ***IL8*** | TTG GCA GCC TTC CTG ATT TC | TGG CAA AAC TGC ACC TTC AC |
| ***IL6*** | CCC TGA GAA AGG AGA CAT GTA AC | CCT CTT TGC TGC TTT CAC ACA TG |
| ***IRF1*** | CTC CAC CTC TGA AGC TAC AAC AG | TCC AGG TTC ATT GAG TAG GTA CC |
| ***IL6ST*** | AAA TCT GAA TGG GCA ACA CAC AAG | CAC AGT AGA ATA ATC AAC AGT GCA TG |
| ***UBC*** | GGG TCG CAG TTC TTG TTT GTG G | GAT GGT GTC ACT GGG CTC AAC |
| ***SOCS5*** | CCG CTA CAA CAG ATC CCT GC | AGT CCC GTT ACA GTG GAG GAG |
| ***JAK2*** | TGG AGC TTT GGA GTG GTT CT | GCA TAA ATT CCG CTG GTG GAC |
| ***IFNGR1*** | CCG AGA TGG AAA AAT TGG ACC AC | CTG AAG GGT GAA ATA TGT CAA TCA TG |
| ***STAT3*** | TTC TTC AGA GCA GGT ATC TTG AGA AG | TCT GTA GAA GGC GTG ATT CTT CC |
| ***JAK3*** | CTC TGG ACT TTG CCA TCA ACA AG | CAC AGA CAG TGA GGA GGA AGC |
| ***ISG15*** | CAT GGG CTG GGA CCT GAC G | TGC ACG CCG ATC TTC TGG G |

**TaqMan Primers and probes**

| **Gene** | **Forward Primer**  **(5' – 3')** | **TaqMan Probe**  **(5' – 3')** | **Reverse Primer**  **(5' – 3')** |
| --- | --- | --- | --- |
| ***MMP1*** | TGT GGA CCA TGC CAT TGA GAA | TCT GCT TGA CCC TCA GAG ACC | AGC CTT CCA ACT CTG GAG TAA TGT CAC ACC |
| ***MMP3*** | GGG CTA TCA GAG GAA ATG AG | CAC GGT TGG AGG GAA ACC TA | AGC TGG ATA CCC AAG AGG CAT CCA CAC |
